# Refined read-out: The hUHRF1 Tandem-Tudor domain prefers binding to histone H3 tails containing K4me1 in the context of H3K9me2/3

**DOI:** 10.1101/2023.07.30.551139

**Authors:** Michel Choudalakis, Goran Kungulovski, Rebekka Mauser, Pavel Bashtrykov, Albert Jeltsch

## Abstract

UHRF1 is an essential chromatin protein required for DNA methylation maintenance, mammalian development and gene regulation. We investigated the Tandem-Tudor domain (TTD) of human UHRF1 that is known to bind H3K9me2/3 histones and is a major driver of UHRF1 localization in cells. We verified binding to H3K9me2/3 but unexpectedly discovered stronger binding to H3 peptides and mononucleosomes containing K9me2/3 with additional K4me1. We investigated the combined binding of TTD to H3K4me1-K9me2/3 *vs*. H3K9me2/3, engineered mutants with specific and differential changes of binding, and discovered a novel read-out mechanism for H3K4me1 in an H3K9me2/3 context that is based on the interaction of R207 with the H3K4me1 methyl group and on counting the H-bond capacity of H3K4. Individual TTD mutants showed up to 10,000-fold preference for the double modified peptides, suggesting that after a conformational change, WT TTD could exhibit similar effects. The frequent appearance of H3K4me1-K9me2 regions demonstrated in our TTD pulldown and ChIP-western blot data suggests that it has specific biological roles. Chromatin pull-down of TTD from HepG2 cells and ChIP-seq data of full-length murine UHRF1 correlate with H3K4me1 profiles indicating that the H3K4me1-K9me2/3 interaction of TTD influences chromatin binding of full-length UHRF1. We demonstrated the H3K4me1-K9me2/3 specific binding of UHRF1-TTD to enhancers and promoters of cell-type specific genes, at the flanks of cell-type specific transcription factor binding sites, and provided evidence supporting an H3K4me1-K9me2/3 dependent and TTD mediated down-regulation of these genes by UHRF1, illustrating the physiological function of UHRF1-TTD binding to H3K4me1-K9me2/3 double marks in a cellular context.

## Introduction

Histone post-translational modifications (PTMs) are a crucial part of chromatin signaling ^1^ with important roles in diseases like cancer ^2^. Among them, histone H3 PTMs have a prominent role with a high number of modifications, some of which are particularly abundant^3^. Over the years, single histone PTMs were found to demarcate various distinct chromatin regions ^1, 4^, for example H3K4me1 marks enhancers ^5^, and H3K4me3 is found on promoters of actively transcribed genes ^6^. In contrast, H3K9me3 is enriched on constitutive heterochromatin^4^ and H3K9me2 occurs in very broad, megabase-long blocks that contribute to inactive chromatin compartment formation ^7^. With a relative abundance exceeding 60%, H3K9me2 is the most common H3 PTM in HeLa cells according to quantitative mass-spectrometry ^8, 9^. In contrast, H3K4 is usually unmodified, and H3K4me1 is the most prevalent modification of this residue with reported abundances of ∼ 30% ^8, 9^. At the next level of complexity, modifications on different residues of H3 were found to co-occur, and ∼ 600 double marks were documented recently ^9^. Synergistic or antagonistic combinations of histone PTMs modulate their biological effects, e.g. enhancers of highly expressed genes harbor H3K4me1 and H3K27 acetylation (H3K27ac) ^5^, while promoters bearing H3K4me3 together with H3K27me3 are silent but poised for activation ^10^. In general, double marks can act through two distinct mechanisms, either by combining the effects of the individual marks, or by signaling new biological outcomes. The latter process depends on ‘reader’ protein domains that are multivalent and bind in a defined manner to multiple marks ^11, 12^.

UHRF1 (Ubiquitin like with PHD And Ring Finger Domains 1) is an E3 ubiquitin ligase with five domains, a Ubiquitin-Like domain (UBL), Tandem-Tudor domain (TTD), Plant Homeodomain (PHD), SET- and RING-associated (SRA) domain, and Really Interesting New Gene (RING) domain (Figure 1A). The TTD was found to bind to H3K9me2/3 ^13, 14^ and microscopy studies showed that TTD is the principal driver of UHRF1 subnuclear localization on heterochromatic H3K9me2/3 foci ^13^. Structural studies showed that the two Tudor domains of TTD form a groove between them, wherein H3 (residues 1-11) peptides containing H3K9me3 bind and place the K9me3 in a classical aromatic cage formed by the TTD residues F152, Y188, and Y191 ^13, 14^. H3K4me2/3 was found to reduce binding in the H3K9me3 context ^13^, while H3S10ph did not ^14^. Further investigations of peptide binding by TTD have revealed many, complex interactions both within the protein and with other partners, for example two autoinhibitory peptides from other parts of the protein can either occupy the H3 binding groove or be allosterically displaced ^15–18^. Moreover, stronger binding of TTD to LIG1-K126me2/3 than to H3K9me2/3 was discovered ^18, 19^. Additional studies showed that PHD can recognize the unmodified H3R2, and the linked TTD-PHD domains were observed to engage a multivalent H3-tail interaction binding H3R2me0-K9me2/3 ^15, 20–22^ connecting UHRF1 to euchromatin ^23, 24^. However, the biological relevance of many of these *in vitro* observations is still unclear ^25^.

**Figure 1.**
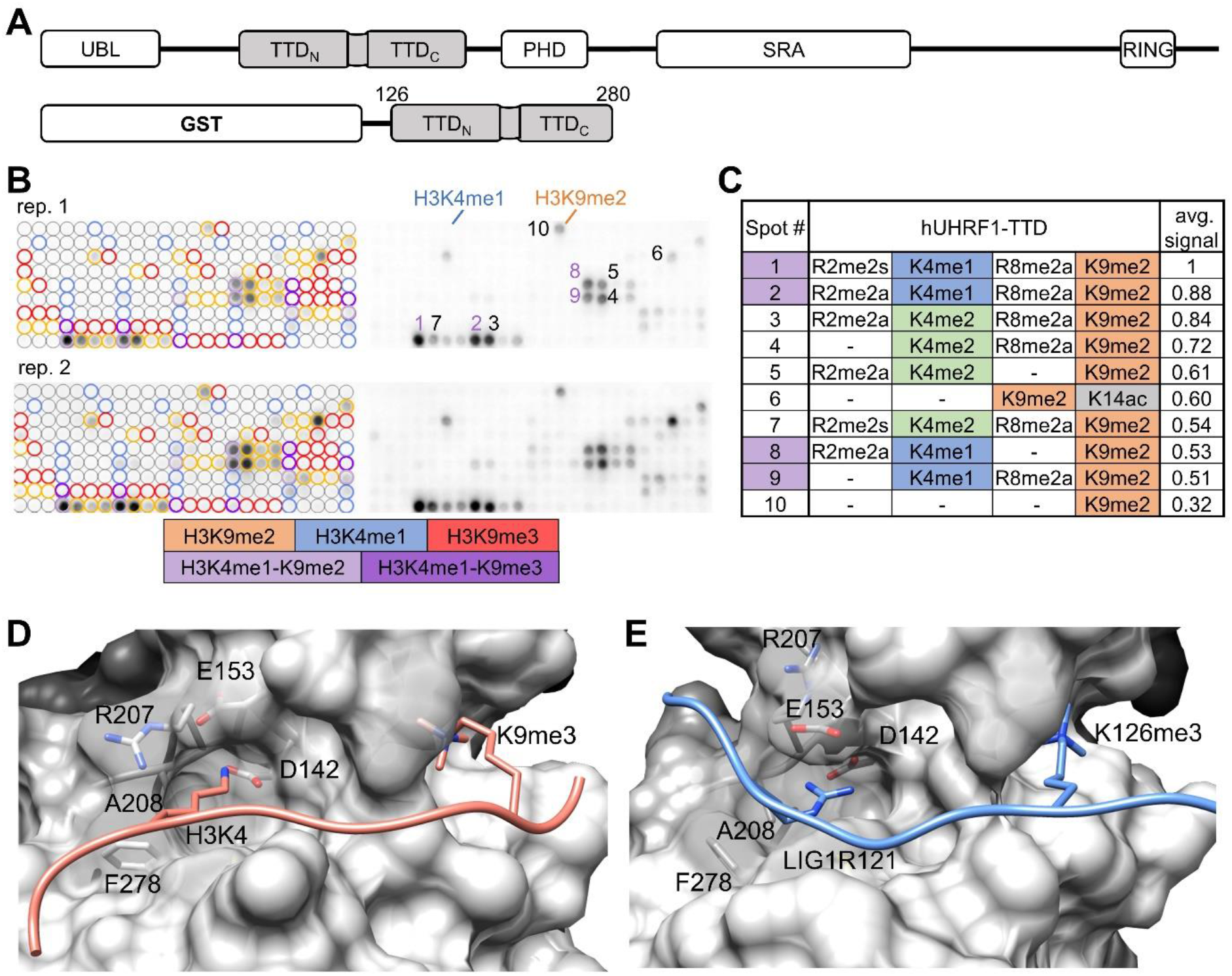
hUHRF1-TTD binds H3 peptides with K4me1 and K9me2 on peptide arrays. **A** Domain structure of the UHRF1 protein containing a Ubiquitin-Like domain (UBL), a Tandem-Tudor domain (TTD), a Plant Homeodomain (PHD), a SET- and RING-associated (SRA), and a Really Interesting New Gene (RING) domain. Bottom: Scheme of the human TTD construct (Uniprot Q96T88, residues 126-280) used here with N-terminal GST-tag. **B** TTD binds to H3K4me1-K9me2 and other H3K9me2 peptides on CelluSpots peptide arrays. Colored circles annotate all peptide spots carrying the selected modifications. The ten best-bound peptides are annotated by order of decreasing average signal. The image shows two independent repeats of the array binding experiment, each containing two technical repeats. **C** Table of the ten best-bound peptides shown in panel (B) and their modifications arranged by decreasing average signal. See also Supplemental Figures 1A and 3. **D** Solution structure of a H3K9me3 peptide - TTD complex (PDB: 2L3R) showing the H3K9me3 peptide bound in an extended conformation in a groove between both Tudor domains with H3K4me0 placed in an acidic pocket (D142, E153) and H3K9me3 in an aromatic pocket. The H3 peptide is shown in light red, TTD in grey surface. **E** Crystal structure of a LIG1K126me3 peptide - TTD complex (PDB: 5YY9), showing that TTD interacts with different peptides in discrete binding modes. The LIG1 peptide is shown in light blue, TTD in grey surface. Panels (D) and (E) were generated with Chimera v1.14 (rbvi.ucsf.edu/chimera). See also Supplemental Figure 1B and 1C. See also Supplemental Figures 1 and 2.

UHRF1 functions as an epigenetic hub, coordinating and recruiting different chromatin interacting proteins ^26, 27^. In 2007, landmark studies revealed the necessity of UHRF1 for DNA replication and maintenance of DNA methylation ^28, 29^. Knockout of UHRF1 in mice is embryonically lethal and UHRF1 KO ES cells show massive DNA hypomethylation, particularly in heterochromatic regions and retrotransposon elements ^28, 30^. Later studies showed that this effect is mediated by UHRF1-catalyzed ubiquitination of H3 that contributes to DNMT1 recruitment ^31^ and by direct interaction of UHRF1 and DNMT1 ^32, 33^. The subnuclear localization of UHRF1 with H3K9me2/3 dependent enrichment in pericentric heterochromatin of interphase nuclei was also found to direct maintenance DNA methylation to these regions ^14^. In recent years, many more processes directly involving UHRF1 have been uncovered, including DNA damage repair, regulation of differentiation and gene regulation ^27^. In cancer cells, UHRF1 is often up-regulated, and it can bind gene promoters to mediate silencing of the associated genes ^27^. Generally, UHRF1 disruption results in strong DNA hypomethylation and reduced cell-type specific gene expression, pointing toward its important role as a regulator of cell lineage specification during differentiation ^34–38^.

In this study, we investigated the human UHRF1-Tandem Tudor domain that binds H3K9me2/3 histones, and is one of the major drivers of UHRF1 localization in cells. We discovered preferential binding of TTD to H3 peptides and mononucleosomes containing K9me2/3 together with K4me1. We describe a novel readout mechanism for H3K4me1 in an H3K9me2/3 context which is based on the interaction of R207 with the H3K4me1 methyl group and on counting the H-bond capacity of H3K4. Interestingly, TTD mutants showed up to 10,000-fold specificity for the double modified peptides, suggesting that specific conformations of TTD exist, which mediate strong H3K4me1-K9me2/3 readout. We demonstrate that TTD specifically binds enhancers and promoters of cell-type specific genes, at the flanks of cell-type specific transcription factor binding sites. The physiological relevance of our findings is demonstrated by showing that published full-length murine UHRF1 ChIP-seq data strongly correlate with H3K4me1 profiles in regions containing H3K9me2/3, indicating that the H3K4me1-K9me2/3 readout by TTD is of key relevance for chromatin binding of UHRF1 in cells. Moreover, by reanalysis of published date we demonstrate H3K4me1-K9me2/3 dependent down-regulation of these genes by UHRF1 that is mediated by the TTD chromatin interaction.

## Results and Discussion

### hUHRF1 Tandem-Tudor preferentially binds H3 peptides with K4me1 and K9me2 on peptide arrays

We purified the hUHRF1-TTD (residues 126-280) ^14^ fused to glutathione S-transferase (GST) (Figure 1A). To screen for combined binding of TTD to H3 peptides with multiple modifications, we used CelluSpots^TM^ peptide arrays ^39^, which contain 275 different H3 histone tail peptides with up to four modifications (Figure 1B). Analysis of the results generated a binding specificity profile of TTD to modified H3 peptides which is shown in Figure 1C. As previously reported ^13, 14^, TTD bound H3K9 methylated peptides, but with a clear preference for H3K9me2 on CelluSpots^TM^ arrays. Unexpectedly, the strongest binding was detected with peptides containing the H3K4me1-K9me2 double modification, while peptides carrying each of the single modifications separately showed no (H3K4me1) or weaker (H3K9me2) binding signals. This indicated a stimulation of binding, resulting from the presence of both modifications on the same peptide. This observation was reproduced in two independent replicates of the experiment (Figure 1C, Supplemental Figure 1A). As the second-best double modification, H3K4me2-K9me2 modified peptides were identified, but not further analyzed. On the CelluSpots arrays, we also observed R-methylation on the preferentially bound H3 peptides (Figure 1C), but unfortunately, the commercial array is lacking H3 peptides only methylated on K4 and K9 without R-methylation. However, two different H3R-methylation sites (R2 and R8) and structural isomers (R2me2a and R2me2s) were observed, indicating the absence of a position and modification specific stimulatory effect, and a stimulatory role of R-methylation on TTD binding was ruled out by additional experiments (Supplemental Figure 2).

Previously, the TTD was shown to bind to various methylated and unmethylated peptides, with structural evidence of changes in binding modes and conformational rearrangements ^40^. In the solution structure of TTD with a bound H3K9me3 peptide, the peptide had an extended conformation lying inside the binding groove between the two Tudor domains with H3K4 placed in an acidic pocket (D142, E153) and H3K9me3 in the aromatic cage formed by F152, Y188 and Y191 (Figure 1D, PDB: 2L3R) ^13^. The LIG1-K126me3 peptide bound to the same groove in a similar conformation, but the acidic pocket was occupied by LIG1-R121, and the aromatic cage bound LIG1-K126me3 (Figure 1E, PDB: 5YY9) ^18^. Strikingly, binding to the LIG1 peptide was more than 100-fold stronger compared to H3K9me3 (9 nM *vs*. 1600 nM). This effect was due to the binding of LIG1-R121 into the acidic pocket, as seen by the strongly elevated binding strength of an K4R-K9me3 H3 mutant peptide (22 nM) ^18^, and it could be attributed to the better geometry of the bidentate H-bonds between UHRF1-D142 and LIG1-R121. Another characteristic of the different binding mode was the lack of interaction of LIG1-R121 with UHRF1-E153 in the H3K4 binding pocket (Supplemental Figure 1B-C). These data suggest that the acidic binding pocket is not ideally occupied with an unmodified lysine residue, which is relevant in the context of the current study, as the preferred binding of the H3K4me1-K9me2 peptide is also expected to be caused by differences in the H3K4me0 *vs*. H3K4me1 interaction in this pocket.

### UHRF1-TTD binds to H3K4me1-K9me2/3 peptides better than to H3K9me2/3

To validate our peptide array binding results, we conducted equilibrium peptide binding experiments using fluorescently labelled H3 peptides with the single K9me2/3 or double K4me1-K9me2/3 modifications (Supplemental Table 2). In fluorescence anisotropy (FA) titrations with GST-fused TTD, we determined the TTD dissociation constant (K_D_) of H3K9me3 to be 680 ±18 nM but 240 ±51 nM for H3K4me1-K9me3 (Figure 2A). This corresponds to an approximately 3-fold stimulation of binding by H3K4me1 being present together with H3K9me3 on the same peptide (Table 1). Studies with the H3K9me2 and H3K4me1-K9me2 peptides showed a similar trend, with a >2-fold preference for the double modified peptide (Supplemental Figure 3A). The measured K_D_ values for H3K9me2/3 binding agree with the literature ^14, 15, 22^. Control experiments with H3 peptides without modification or carrying only K4me1 confirmed the necessity of K9me2/3 for strong interaction (Supplemental Figures 3A and 3B). Thus, we validated and quantified the preferential binding of TTD to H3K4me1-K9me2/3 peptides compared to H3K9me2/3. To the best of our knowledge, the only previously reported binding titrations with a H3K4me-K9me peptide were with H3K4me3-K9me3, showing a weakened interaction with TTD ^13^.

**Figure 2.**
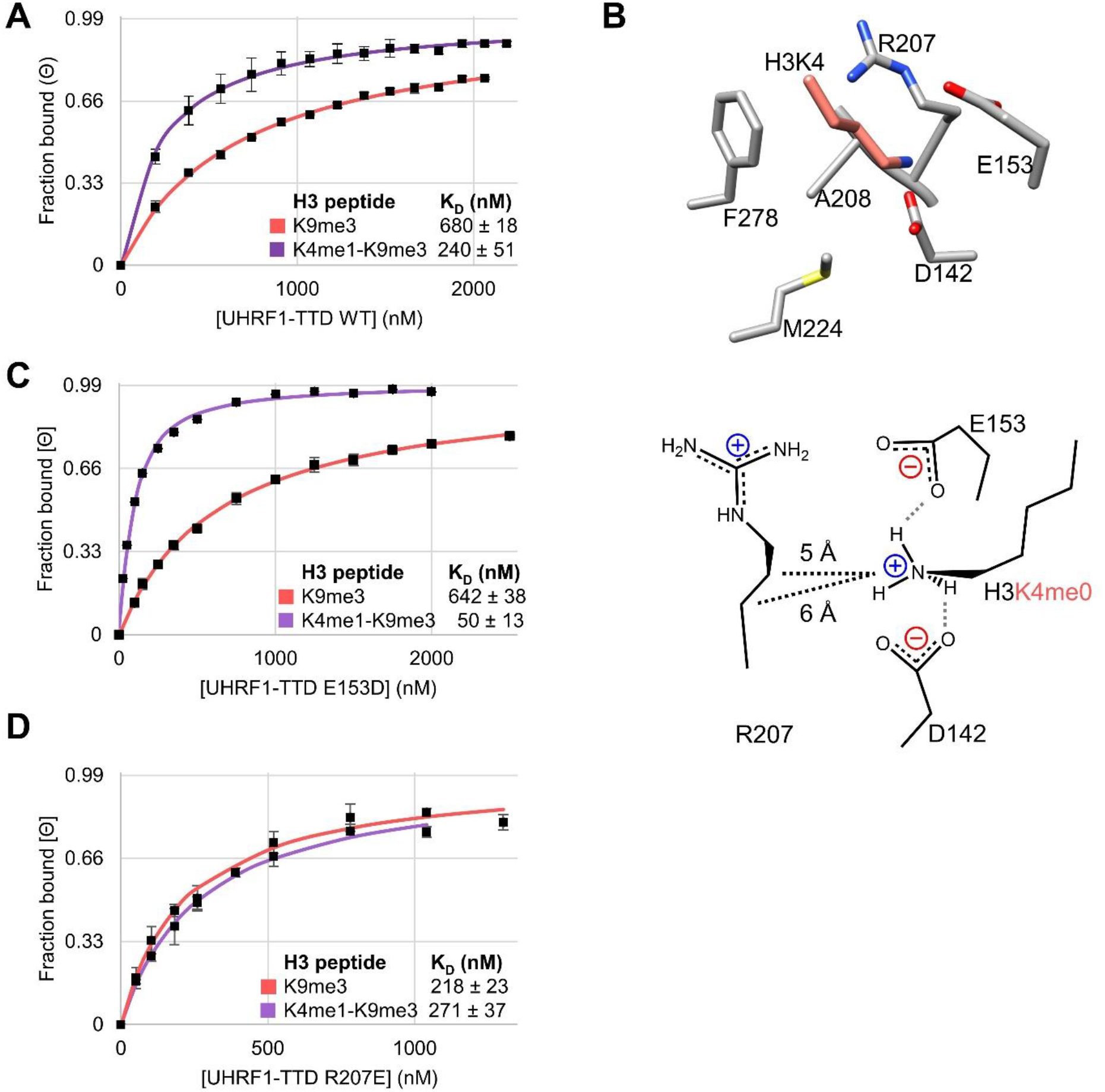
hUHRF1-TTD binds to H3K4me1-K9me3 peptides better than to H3K9me3 alone and adopts discrete binding modes. **A** TTD wild-type (WT) binds H3K4me1-K9me3 peptides more strongly than H3K9me3 in equilibrium peptide binding titrations analysed by the fluorescence anisotropy (FA) change. Data points are average fraction bound (Θ) of *n* ≥2 independent experiments, error bars are 0.95 confidence intervals (CI). K_D_ are the average of *n* ≥2 independent fits, errors are 0.95 CI. **B** Snapshot of the H3K4 binding pocket in the H3 – TTD complex (PDB: 2L3R) showing the investigated residues. Scheme: Model of the interactions in the H3K4 binding pocket for H3K4me0 readout in H3K9me3 context. H-bonds in gray dashed lines, van der Waals contacts in black dashed lines. **C** Representative data showing that TTD E153D binds H3K4me1-K9me3 peptides more strongly than WT. **D** Representative data showing that TTD R207E binds H3K9me3 peptides more strongly than WT. See also Supplemental Figure 3.

**Table 1.** Equilibrium peptide binding of hUHRF1-TTD with H3(1-19) peptides. Binding constants were detected by fluorescence anisotropy using FITC labeled peptides. Dissociation constants (K_D_) are reported as the mean and 0.95 confidence intervals (CI) of *n* ≥ 2 independent titrations. Relative effect of mutation is the ratio of K_D_ for WT over K_D_ for Mut, and >1 signifies a stronger binding for that mutant with this specific peptide. FITC - fluorescein isothiocyanate; WT – wild type; Mut – mutant.

| UHRF-TTD | Average $K_D \pm 0.95$ CI (nM) | | | Rel. effect of mutation<br>($K_D$ WT/ $K_D$ Mut) | |
| --- | --- | --- | --- | --- | --- |
|  | H3<br>K9me3 | H3<br>K4me1-K9me3 | Ratio | H3<br>K9me3 | H3<br>K4me1-K9me3 |
| WT | 680 ($\pm 18$) | 240 ($\pm 50$) | 2.8 | - | - |
| D142A | 4900 ($\pm 300$) | 2700 ($\pm 220$) | 1.8 | 0.14 | 0.09 |
| D142E | 980 ($\pm 120$) | 580 ($\pm 4$) | 1.7 | 0.69 | 0.41 |
| D142N | 4500 ($\pm 200$) | 2 ($\pm 0.1$) | 2800 | 0.15 | 150 |
| E153A | 16000 ($\pm 490$) | 2 ($\pm 1$) | 9600 | 0.04 | 140 |
| E153D | 640 ($\pm 40$) | 50 ($\pm 19$) | 13 | 1.1 | 4.8 |
| E153Q | 13000 ($\pm 4600$) | 250 ($\pm 55$) | 51 | 0.05 | 0.95 |
| R207A | 900 ($\pm 110$) | 370 ($\pm 52$) | 2.5 | 0.74 | 0.65 |
| R207L | 900 ($\pm 130$) | 440 ($\pm 14$) | 2 | 0.77 | 0.54 |
| R207Q | 1600 ($\pm 230$) | 390 ($\pm 70$) | 4.1 | 0.42 | 0.62 |
| R207H | 9700 ($\pm 1150$) | 460 ($\pm 10$) | 21 | 0.07 | 0.52 |
| R207E | 220 ($\pm 20$) | 270 ($\pm 36$) | 0.8 | 3.1 | 0.9 |
| A208G | 9500 ($\pm 1600$) | 2400 ($\pm 180$) | 4 | 0.07 | 0.1 |
| M224A | 1400 ($\pm 270$) | 430 ($\pm 45$) | 3.3 | 0.48 | 0.55 |
| F278A | 7900 ($\pm 1600$) | 3900 ($\pm 710$) | 2 | 0.09 | 0.06 |

Next, we were curious to understand the mechanism of the stimulation of peptide binding by H3K4me1 in the context of H3K9me3. Taking a closer look into the H3K4 binding pocket of TTD, D142 and E153 were observed to form H-bonds to K4 (Figure 2B). E153 also interacts with R207 creating a system comprising two interacting acidic and two basic residues. We mutated residues that might interact with the H3K4me1 methyl group to eliminate or weaken the difference between the K_D_ values of H3K9me3 peptides with and without H3K4me1. H3K9me3 peptides were used in these experiments, to allow direct comparison with available TTD-peptide structures. Considering aromatic-hydrophobic and hydrophobic-hydrophobic interactions, we selected the UHRF1-TTD mutants F278A, M224A and A208G for analysis (Table 1, Supplemental Figure 3C). In each case, very modest changes of the preference towards the double modified substrate were observed suggesting these residues were not involved in the H3K4me1 readout. Next, we mutated D142 to A and E, and both showed similar preference ratios, but D142A showed a strongly reduced binding of both peptides (Supplemental Figures 3D and 3E). However, with the D142N mutation, the preference for binding the double modified peptide was strongly elevated, indicating that the two peptides interacted differently with this mutant (Table 1). Next, the role of E153 was investigated. E153D bound the H3K9me3 peptide similarly as WT (Figure 2C), but with H3K4me1-K9me3 we observed a gain in binding. E153A/Q led to a near-complete loss of the H3K9me3 interaction (Table 1). Unexpectedly, the H3K4me1-K9me3 peptide was bound with similar to WT or even elevated binding affinity indicating that E153A showed a drastic increase in the preference for binding H3K4me1-K9me3, similar as with D142N. This result demonstrates that both peptides are bound by TTD in distinct conformations and mutations in TTD can trigger its change into a more selective conformation. To investigate if any other residue contributes to the positive binding effect of H3K4me1, we compared the structures of the H3 and LIG1 peptides bound to TTD and observed that R207 had different orientations (Supplemental Figure 1C). Moreover, given the distance of the Nε of H3K4 to the Cβ and Cγ of R207 (6.0 and 4.8 Å, respectively), the H3K4me1 methyl group would be in van der Waals (vdW) contact distances with these methylene groups, explaining the stimulatory effect of H3K4me1 on peptide binding to TTD. We investigated peptide binding of several R207 mutants and observed that R207E lost the preference for binding to H3K4me1-K9me3 (Figure 2D), suggesting that the charge inversion mutation induced a conformational change that disrupted the vdW contact.

The mutant binding data can be compiled leading to a new Kme1 binding mechanism for H3K4me1-K9me3. H3K4me0 has a higher H-bonding potential than H3K4me1 and, therefore, it can interact with both D142 and E153. Consequently, reduction of the H-bonding potential of these residues by D142N, E153A, or E153Q mutations affects K4me0 binding more than K4me1 binding leading to an increased preference for H3K4me1-K9me3. The low binding of H3K4me0-K9me3 and H3K4me1-K9me3 peptides by D142A shows that at least one H-bond from D142 is needed for binding of any of the peptides. The reduced binding of the H3K4me0-K9me3 peptide by D142N and E153A suggests that WT TTD K4me0 forms a bidentate H-bond with D142 and an additional H-bond with E153 (Figure 2B). In contrast, K4me1 can form only two H-bonds, one with D142 and one with an additional H-bond acceptor (either the second oxygen atom of D142 or in its absence E153). In addition, the methyl group of K4me1 makes a vdW contact to R207. This Kme1 interaction mode is distinct from previous models for binding of Kme1/2 which were based on incomplete aromatic cages combined with H-bonds to the amino group ^41, 42^. The important role of H-bonds to the K4 side chain for the TTD interaction can explain the reduced binding of H3K4me3-K9me3 peptides, in which the H-bonding capacity of K4 is fully blocked ^13^. One of the most striking and unexpected results of the mutant analyses was the identification of the E153A TTD mutant showing a 10,000-fold preference for binding to H3K4me1-K9me3 double modified peptides which was due to a strong increase of the binding to double modified peptide and reduced interaction with H3K9me3. This observation suggests that a conformational change moving E153 away from the K4 binding pocket could lead to a similar enhancement of dual mark binding specificity in WT TTD. This hypothesis is in agreement with well documented conformational changes of UHRF1 that demonstrably have important biological outcomes (Fang et al., 2016; Gelato et al., 2014; Kori et al., 2019; Vaughan et al., 2019).

### UHRF1-TTD binds native nucleosomes with both H3K4me1 and H3K9me2/3

We considered the occurrence of the H3K4me1-K9me2/3 double mark in human cells very likely, given the high abundance of the individual PTMs observed in mass spectrometric analyses ^8, 9^. To validate this presumption, H3K4me1 ChIP experiments combined with H3K9me2 western blot detection were carried out. We first validated the specificity of binding for an α-K4me1 ChIP grade antibody under stringent IP conditions and an α-K9me2 antibody was validated under western blot conditions (Supplemental Figure 4A, Supplemental Table 3). Then, using the tested conditions (Supplemental Table 4), we performed H3K4me1 ChIP using mononucleosomes isolated from HepG2 cells followed by H3K9me2 detection using western blot. From the H3 precipitated using the α-K4me1, the α-K9me2 detected robust signals in multiple biological replicates (Supplemental Figure 5A). Relative to input, the α-K4me1 ChIP signals correspond to ∼ 9% of the global H3K9me2 (Supplemental Figure 5B). IgG serves as control for unspecific interactions. These data demonstrate the abundant coexistence of H3K4me1 and H3K9me2 on nucleosomes using antibody-based enrichment and detection.

Having validated the preferential binding of TTD to H3K4me1-K9me2/3 modified peptides, we wanted to examine if the preferential interaction can also be seen with native nucleosomes. To this end, we applied CIDOP (Chromatin Interacting Domain Precipitation) ^43^, an assay similar to chromatin immunoprecipitation (ChIP), using the GST-tagged TTD domain to capture native mononucleosomes isolated from HepG2 cells (Figure 3A), followed by western blot analysis for specific H3 PTMs with ChIP-grade antibodies (Supplemental Table 3), that were carefully validated before use (Supplemental Figure 4B). After optimizing the washing conditions to reduce unspecific and weak interactions, the pull-down of native mononucleosomes with TTD demonstrated enrichment in both H3K4me1 and H3K9me2 (Figure 3B, Supplemental Figure 5C and D). Depletion of H3K4me3 validated the specificity of the assay. To test for unspecific interactions, the same mononucleosome preparation was assayed by CIDOP with the D142A mutant. As positive controls, MPP8-CD (M-Phase Phosphoprotein 8 – Chromo Domain), a known reader of H3K9me2/3 ^39, 44^, and TAF3-PHD (TATA-box binding protein Associated Factor 3 - Plant Homeodomain), an H3K4me3 reader ^45^, were used. The assays were conducted under stringent conditions (Supplemental Table 4) and repeated for a minimum of 3 independent biological replicates showing enrichment of H3K4me1 and H3K9me2 with TTD, but not its D142A mutant, together with enrichment of H3K9me2 with MPP8 and H3K4me3 with TAF3 (Figure 3B, Supplemental Figure 5C and D). As an alternative readout, quantitative PCR (qPCR) was applied to detect the enrichment of H3K9me2 and depletion of H3K4me3 reporter regions in two biological replicates of the TTD CIDOP (Supplemental Figure 6A-C). Control experiments with the binding deficient TTD D142A mutant and IgG demonstrated the specificity of the enrichment, and control ChIP-qPCR/CIDOP-qPCR experiments with α-H3K9me2, MPP8-CD and TAF3-PHD verified the assayed amplicons. In agreement with the western blot results, the qPCR assays revealed enrichment of H3K9me2 and depletion of H3K4me3 with TTD and loss of the H3K9me2 enrichment for the D142A mutant.

**Figure 3.**
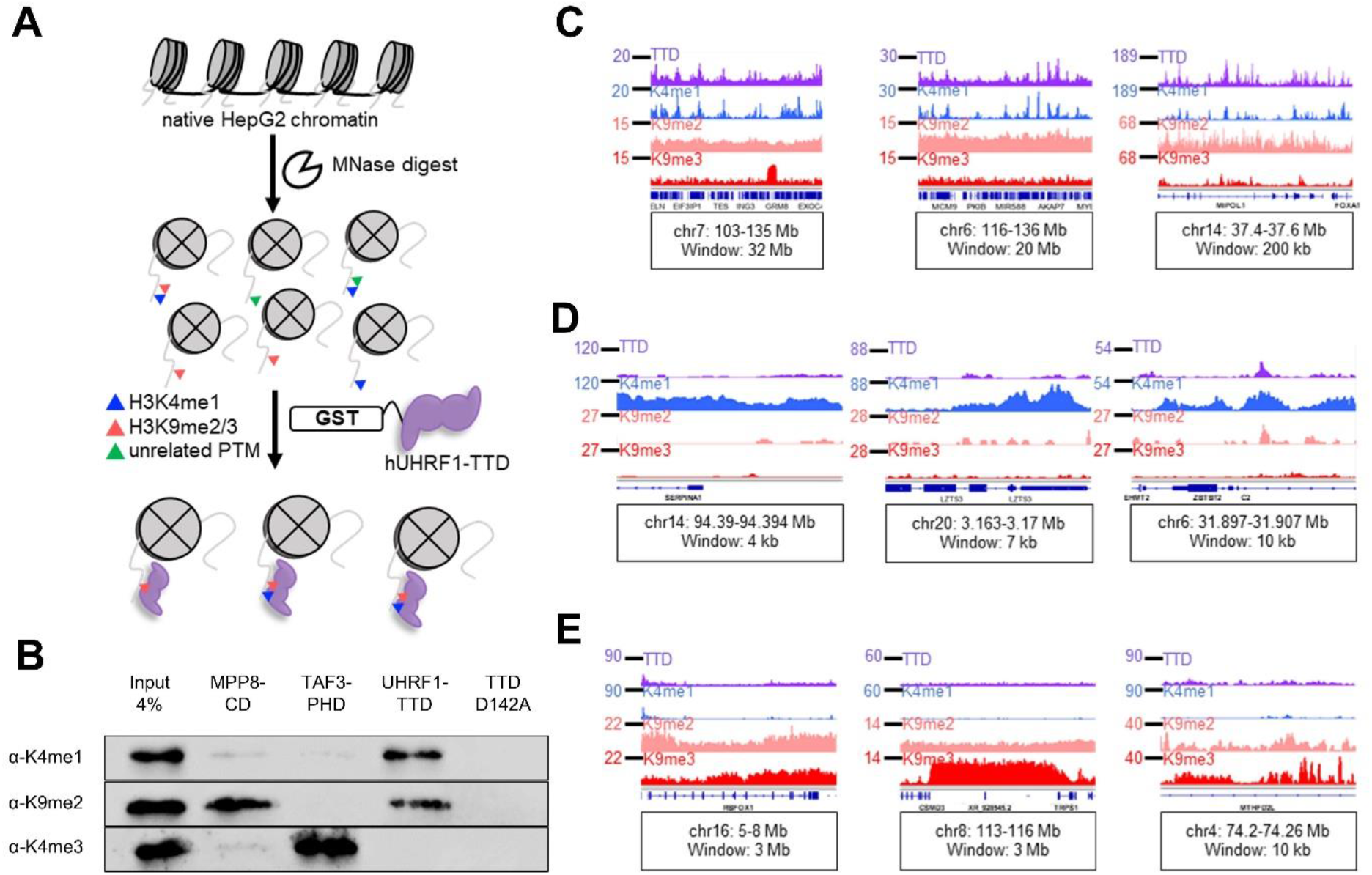
hUHRF1-TTD CIDOP pull-down is enriched in regions with both H3K4me1 and H3K9me2/3. **A** Workflow of Chromatin Interacting Domain Precipitation (CIDOP) experiments used to investigate enrichment of mononucleosomes in characteristic H3 PTMs during pull-down with GST-TTD. **B** TTD pull-down is enriched in H3K4me1 and H3K9me2, but depleted from H3K4me3. Pull-down with the TTD D142A mutant does not show any enrichment. Control experiments with MPP8-CD ^43^ and TAF3-PHD ^45^ showed enrichment of H3K9me2 and H3K4me3, respectively. Shown are representative experiments of *n* ≥3 biological replicates. See also Supplemental Figures 4, 5 and 6. **C** Exemplary browser views showing strong correlation of UHRF1-TTD with H3K4me1 in regions with broad H3K9me2/3 signal. TTD and H3Κ9me2 tracks were derived from two pooled biological replicates. H3Κ4me1 and H3Κ9me3 data were taken from public datasets of comparable HepG2 cells ^47^. See also Supplemental Figures 7 and 8. **D** Exemplary browser views showing lack of TTD enrichment in regions with H3K4me1 but without H3K9me2/3 signal. See also Supplemental Figures 7 and 8. **E** Exemplary browser views showing lack of TTD enrichment in regions with H3K9me3 alone. See also Supplemental Figure 9. All tracks in RPKM, y-axes start from 0. All coordinates in hg38, gene annotation from RefSeq. Browser views of CIDOP-/ChIP-seq data were created with the Integrative Genomics Viewer (software.broadinstitute.org/software/igv/).

To look more deeply into the specific genome-wide binding pattern of TTD, we generated paired-end high-throughput sequencing data from the two biological replicates of the CIDOP reaction with TTD and the H3K9me2 ChIP. The high-quality reads were mapped to hg38, quantified excluding blacklisted regions ^46^, and pooled (Supplemental Figure 7). The genome-wide Pearson’s correlation coefficient (*r*) between the pooled data from UHRF1-TTD CIDOP and each of the replicates was 0.93 and 0.95, respectively. The H3K9me2 ChIP were pooled with *r* 0.93 and 0.86. For comparison, public ChIP-seq data for H3K4me1 and H3K9me3 in HepG2 cells were retrieved ^47^. The genome-wide *r*-values of the pairwise correlation of TTD data with H3K4me1, H3K9me2 and H3K9me3, were 0.58, 0.53 and 0.12, respectively, indicating that TTD does not show a strong correlation with any of the isolated marks, but moderate similarity exists between the TTD, H3K4me1 and H3K9me2 tracks. It should be noted that the colocalization of UHRF1-TTD and H3K9me2/3 at heterochromatic sites ^13, 48^ was not reflected in this analysis, due to the low coverage of heterochromatic fragments in the ChIP-seq data. For further analysis, we visualized the TTD data alongside H3K4me1, H3K9me2, and H3K9me3, seeing a very strong correlation of TTD binding with H3K4me1 profiles in areas with broad H3K9me2/3 signal (Figure 3C). This unexpected observation strongly supports our previous biochemical data revealing a preferred binding of TTD to H3K4me1-K9me2/3 double marks. Regions rich in H3K4me1, but with little H3K9me2/3, showed negligible TTD signals (Figure 3D). At the same time, regions with H3K9me2/3 alone showed low TTD signal (Figure 3E), indicating a conditional contribution of the H3K4me1-K9me2/3 double mark for robust TTD binding. The colocalization of TTD, H3K4me1, and H3K9me2 and the resemblance of peak motifs was seen repeatedly at different genomic regions (Supplemental Figure 8). These findings indicate that H3K4me1 and H3K9me2/3 are the principal marks behind the TTD signal in non-repetitive genomic regions. Additionally, we verified the broad megabase-wide distributions of H3K9me2 lacking defined peaks ^7^ (Supplemental Figure 9). Jointly, the genome-wide analysis by western blot and NGS confirmed the specific enrichment of UHRF1-TTD pull-down in nucleosomes carrying H3K9me2 as well as H3K4me1.

### UHRF1-TTD CIDOP prefers native H3 with both K4me1 and K9me2/3

Bringing together our *in vitro*, western blot, and NGS data so far, we hypothesized that TTD binding occurs in regions with broad distribution of H3K9me2, in which H3K4me1 peaks resulted in stronger TTD interaction. Based on this, we expected stronger TTD signal in regions enriched in H3K9me2, where strong TTD signals should correlate with strong H3K4me1. To test for this, we divided the entire genome in 1 kb bins, arranged them by decreasing mean H3K9me2 signal in 10 equal deciles as done in Ming et al. 2020 ^49^. For each, we plotted the mean signals of TTD (Figure 4A), H3K4me1, and H3K9me2 and observed that TTD signals followed the decreasing trend of H3K9me2 across deciles, unlike H3K4me1 (Supplemental Figure 10A). Looking within decile 1, the gradients of H3K9me2 and TTD are similar (Figure 4B). Furthermore, in H3K9me2 rich deciles, the TTD signal showed a high correlation with H3K4me1 (*r* 0.8) that declined in deciles with lower H3K9me2 signal (Figure 4C). This demonstrated that a strong TTD signal is observed at regions with a robust, broad H3K9me2 signal, and within these H3K4me1 modulates TTD intensity. As an additional control, we plotted a heatmap of TTD, H3K4me1, H3K9me2 and H3K9me3 centered on H3K4me1 peaks (Figure 4D), addressing the abundance of the H3K4me1-K9me2/3 double mark and showing that about 2/3 of the H3K4me1 peaks also contain H3K9me2. This finding indicates that the H3K4me1-K9me2 double mark frequently occurs in the non-repetitive part of the genome. As there is little information about H3K4me1 in heterochromatin, no statements about the co-occurrence of H3K4me1-K9me2/3 in this part of the genome can be made. As expected from the previous analyses, the intensity of the TTD signal was better correlated to H3K9me2 than to H3K4me1 (*r* 0.76 *vs.* 0.25), because not all H3K4me1 peaks carry H3K9me2 and the similarity in patterning between TTD and H3K4me1 was only evident in regions with robust H3K9me2.

**Figure 4.**
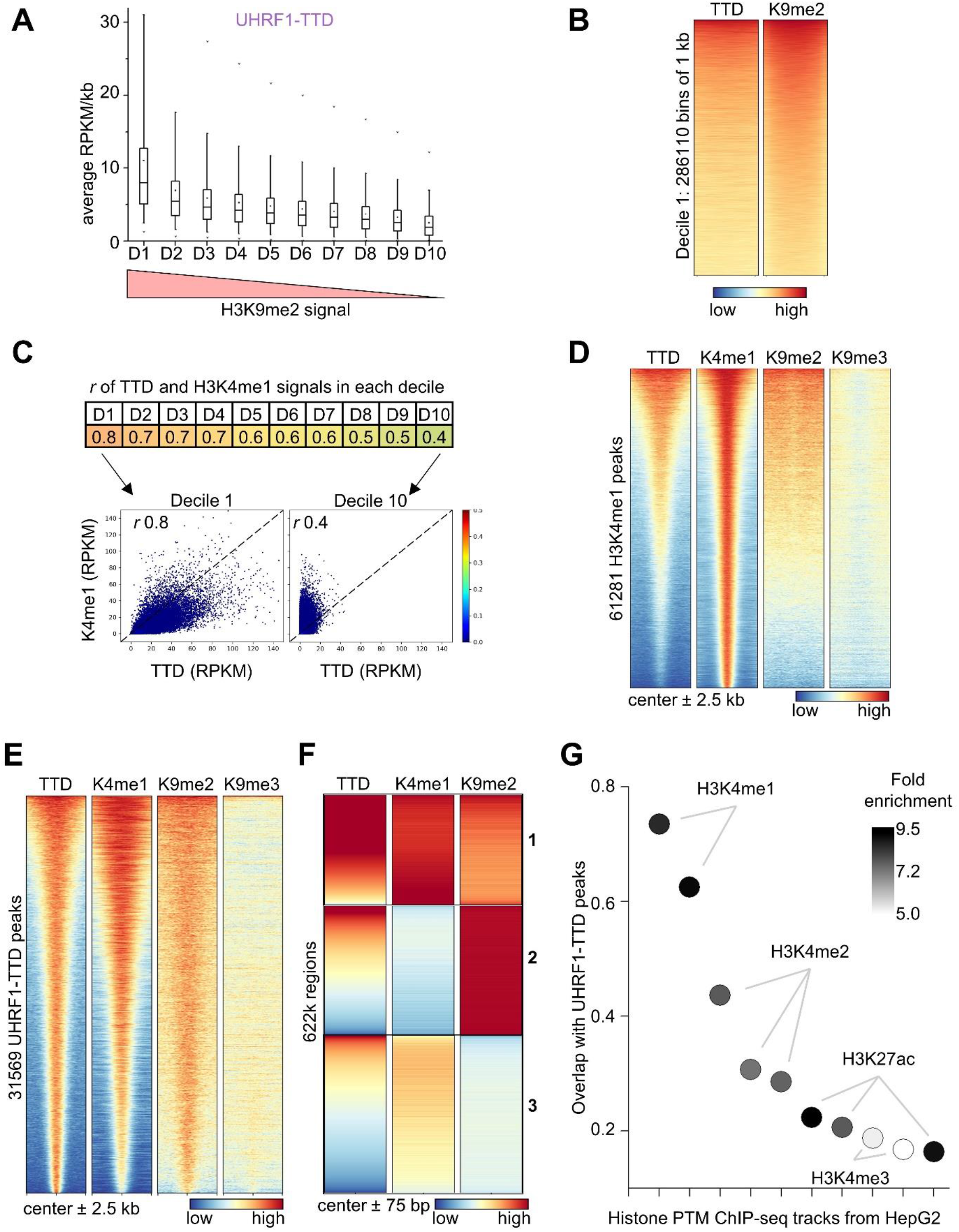
hUHRF1-TTD shows strong correlation with H3K4me1 at H3K9me2 regions. **A** TTD signal is strongest in H3K9me2 highly enriched genomic regions and follows the decreasing H3K9me2 signal strength. The entire genome was divided into 1 kb bins, arranged by decreasing mean H3K9me2 signal, divided into deciles and the mean TTD signal of each group was plotted. Central line is median, box borders are 25^th^ to 75^th^ percentile, and whiskers 5^th^ to 95^th^. See also Supplemental Figure 10A. **B** Heatmap of decile 1, showing regions of genome-wide highest H3K9me2 signal and the corresponding TTD signal. Heatmap of 286110 bins of 1 kb, arranged by decreasing H3K9me2 signal. **C** TTD to H3K4me1 correlation is strongest in genomic regions with high H3K9me2. *r*-values of TTD and H3K4me1 signals were calculated for the deciles shown in panel (A). Within regions with high H3K9me2, average TTD signal has high correlation with H3K4me1, which declines as H3K9me2 signal decreases. **D** H3K4me1 peaks contain H3K9me2 signal. Within regions with H3K9me2 and H3K4me1, TTD shows enrichment. Heatmap of 61281 K4me1 peaks and their ± 2.5 kb flanks, centered in the middle, arranged by decreasing signal. **E** TTD peaks contain H3K4me1 peaks and enriched H3K9me2 signal. Heatmap of all 31569 TTD peaks and their ± 2.5 kb flanks, centered in the middle, and arranged by decreasing TTD signal. See also Supplemental Figure 10B. **F** Heatmap of TTD peaks split into ∼ 622k 150 bp-wide fragments, clustered by *k*-means, and arranged by decreasing TTD signal. The strongest signal inside TTD peaks comes from mononucleosomes with H3K4me1 and H3K9me2. See also Supplemental Figure 10C. **G** Overlap of TTD peaks with public ChIP-seq peaks, individually curated to only include comparable HepG2 cells. Data shown here are the first ten ChIP-seq datasets with the highest overlap in TTD peaks in the public ChIP-Atlas database (chip-atlas.org) ^50^, as arranged by decreasing percentage of TTD peak overlap (counts). Each has log *p* < -10. Circle shading reflects the enrichment over randomized input.

To further validate our finding of the combined readout of H3K4me1 and H3K9me2/3 by TTD, we also performed broad peak calling on TTD. Due to its very broad distribution, peak calling was not possible on H3K9me2, but broad peaks could be identified on H3K9me3. Using the broad peaks of the TTD enrichment, we prepared a heatmap of TTD, H3K4me1, H3K9me2 and H3K9me3 signals centered on these regions. The data clearly showed that TTD peaks have a strong enrichment and positive correlation with H3K4me1 (Figure 4E), and 2/3 of the TTD peaks overlap with H3K4me1 peaks (Supplemental Figure 10B). At the same time, TTD peaks also correlated with a gradient of H3K9me2, with little detectable contribution from H3K9me3, finally revealing a clear similarity of patterning for TTD, H3K4me1 and H3K9me2. This validated that the strong UHRF1-TTD pull-down signal originated from a colocalization of H3K9me2 and H3K4me1, with a conditional contribution from each mark, suggestive of combined TTD binding. To further validate the preferential binding of UHRF1-TTD to mononucleosomes with double H3K4me1-K9me2/3 marks, the broad TTD peaks were split into ∼ 622k mononucleosome sized 150 bp fragments which were then used in *k*-means clustering, resulting in 3 clusters that were arranged by decreasing TTD signal (Figure 4F). This analysis confirmed that, within the broad UHRF1-TTD CIDOP peaks, the strongest TTD signal came from fragments bearing H3K4me1-K9me2 (cluster 1), followed by regions rich in H3K9me2 some of which showed strong TTD signals (cluster 2), while the weakest signal was found in regions with H3K4me1 but with low amounts of H3K9me2 (cluster 3). As these signals are based on the enrichment of 150 bp DNA fragments, they clearly indicate the presence of both marks on one mononucleosome. Addition of H3K9me3 data to this analysis revealed that TTD and H3K9me3 signals were correlated in cluster 3 (Supplemental Figure 10C) indicating that in this cluster combined readout of H3K4me1-K9me3 occurred.

To better address TTD binding to H3K4me1-K9me3 without K9me2, we clustered the H3K9me3 peaks and arranged the clusters by increasing H3K9me2 signal (Supplemental Figure 11A). In one cluster of this analysis (#2), we clearly observed TTD binding to H3K4me1 and H3K9me3 in absence of H3K9me2. To validate this finding in reverse, H3K9me3 peaks were found on 12% of H3K4me1 peaks (Supplemental Figure 11B). A heatmap of H3K4me1 peaks overlapping to ≥50% H3K9me3 peaks and clustering again revealed one cluster (#2), where TTD binds to H3K4me1 and H3K9me3 in absence of H3K9me2 (Supplemental Figure 11C). Hence, while the biologically more abundant double modified substrate in the non-repetitive loci of HepG2 cells is H3K4me1-K9me2, our *in vitro* and NGS data both document the preferential binding of TTD to H3K4me1-K9me2 and H3K4me1-K9me3.

For an independent validation of H3K4me1 as the second part of the double mark read by TTD, we used the TTD peaks and conducted peak overlap analysis with various public ChIP-seq data deposited in the ChIP-Atlas database (chip-atlas.org) ^50^. The results were restricted to ChIP-seq experiments in HepG2 wild-type cells analyzing abundant PTMs and arranged by decreasing TTD peak overlap. The first ten ChIP-seq datasets with the highest overlap in TTD were plotted and analyzed for fold enrichment of ChIP-seq peaks in TTD peaks over randomized controls (Figure 4G and Supplemental File 1). Strikingly, two H3K4me1 tracks from independent laboratories had the highest overall overlap with TTD peaks (73% and 63%, respectively) and more than 8-fold enrichment. H3K4me2 tracks completed the top-five, but showed a smaller overlap of 44% or less, and less significant enrichment. This clearly demonstrates that among all the publicly available datasets for histone ChIP-Seq in HepG2 cells, H3K4me1 peaks have the best correlation to TTD peaks. Unfortunately, no public H3K9me2 ChIP-seq data were found for HepG2 that could be used as independent validation of our own data.

In summary, our data demonstrate that binding of GST-tagged UHRF1-TTD to native HepG2 chromatin required H3K9me2/3, with higher affinity for mononucleosomes with double H3K4me1-K9me2/3 modifications, supporting our hypothesis of combined binding of both marks and complementing our peptide-binding data. Moreover, we document the wide occurrence of the H3K4me1-K9me2 double mark in TTD CIDOP and ChIP-western blot experiments.

### UHRF1-TTD binds on promoters of cell type specific genes and down-regulated genes in HepG2

Having established the preferential enrichment of TTD CIDOP in H3K4me1-K9me2 regions from HepG2 chromatin, we wondered which functional role could be attributed to TTD binding to the H3K4me1-K9me2 double mark. It is well known that H3K4me1 marks enhancers, and H3K9me2 has a high abundance and broad distribution (Supplementary Figure 9B). Using the HepG2-specific 18-state ChromHMM reference data (egg2.wustl.edu/roadmap) ^51^, we analyzed the genome-wide distribution of TTD peaks in the different chromatin regions and compared it to randomized controls of equal number and length of peaks (Figure 5A). The enrichment of TTD peaks was high in ‘Enhancers’ (lowest in bivalent enhancers), as well as the regions ‘Flanking TSS’ (upstream or downstream), which include gene promoters. This complements our finding that TTD peaks showed significant overlap with H3K4me1 again indicating that K4 methylation has a marked influence on chromatin binding of TTD. Due to the enrichment of TTD peaks in transcriptional start site (TSS) flanking regions, we looked at all the TSSs from the human TSS reference set (refTSS) ^52^ and since UHRF1 participates in DNA methylation, we also included whole genome bisulfite sequencing (WGBS) data ^53^. Using *k*-means clustering, we identified 4 clusters that were arranged by decreasing TTD signal (Figure 5B). This heatmap shows all human TSSs with their promoter regions, and in clusters 1 and 2, strong or moderate TTD density was observed, respectively. As expected for promoter regions, WGBS DNA methylation levels were relatively low. Interestingly, in clusters 1 and 2, TTD enrichment corresponds to the WGBS signal, both signals showing minima in the center of the regions (at the position of the TSS) and increase towards the flanks. Conversely, in cluster 3 both TTD and WGBS signals are lower. Clusters 3 and 4 function as control regions, the former containing the unmethylated TSSs and promoters of HepG2 cells, while the latter has most of the TSSs with the highest DNA methylation. The patterns observed in clusters 1 and 2 suggest that TTD binding to the regions flanking TSSs can increase DNA methylation at these sites, in agreement with the well-documented role of UHRF1 in the deposition of DNA methylation ^28, 29^.

**Figure 5.**
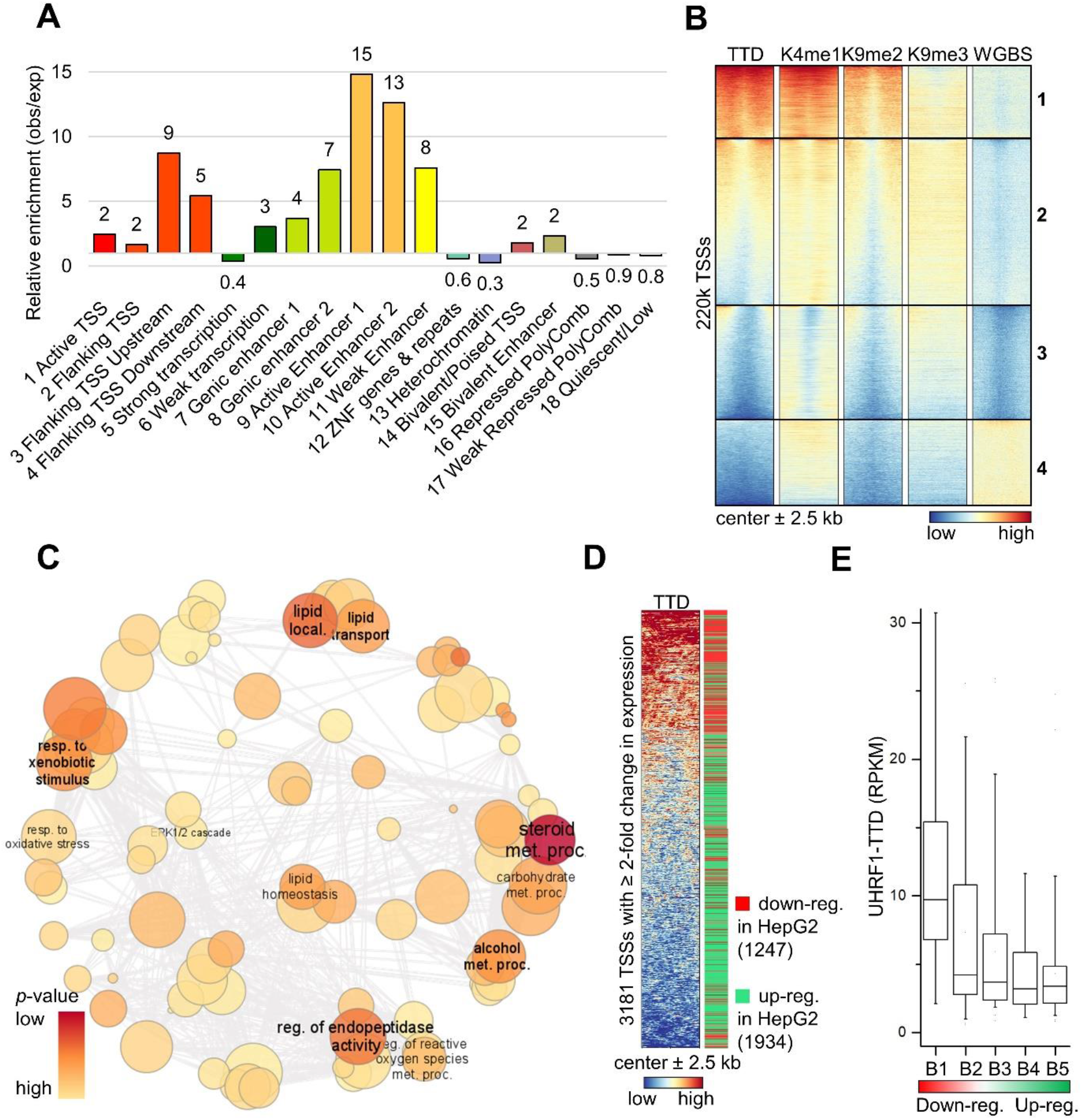
hUHRF1-TTD binds to promoters of cell-type specific genes and down-regulated genes in HepG2. **A** Analysis of TTD peaks in functional chromatin regions. Segmentation data from ChromHMM for HepG2 (egg2.wustl.edu/roadmap) ^51^, enrichment is over randomized control. TTD peaks show strong enrichment in enhancers and TSS flanking regions. **B** Heatmap of ∼ 220k refTSS (centered in the middle ± 2.5 kb) clustered by *k*-means, and arranged by decreasing TTD signal. The TTD enrichment follows H3K4me1 and H3K9me2 signal intensity. WGBS - whole genome bisulfite sequencing. **C** TTD signal is enriched in cluster 1 of panel (B). The genes of this cluster have statistically significant relation to cell-type specific processes for HepG2. refTSS regions were assigned to genes by *ChIP-Enrich* (chip-enrich.med.umich.edu) ^85^, the enriched GO:BP genesets with FDR ≤ 0.05, hybrid p-value ≤ 0.05 were summarized in *Revigo* (revigo.irb.hr) ^86^ and visualized using *Cytoscape* (cytoscape.org) ^87^. Exemplary GO terms for each cluster are annotated. Semantic similarity is reflected in the clustering, color and font-size reflect *p*-value. Circle radius reflects the log of number of genes in GO term ID. dev - development; diff - differentiation; epith - epithelial; local - localization; met - metabolic; pos - positive; proc - processes; reg - regulation; res. – response; stim. – stimulus; trans - transport. See also Supplemental Figure 12A. **D** Heatmap of 3181 refTSS (centered in the middle ± 2.5 kb) corresponding to genes with ≥ 2-fold change in expression between HepG2 and liver tissue and arranged by decreasing TTD signal. The bar on the right is red for down-regulated and green for up-regulated genes in HepG2. Expression data for HepG2 ^84^ and liver cells ^83^ were obtained from The Human Protein Atlas version 21.1 (proteinatlas.org). The TSSs with TTD-rich flanks corresponded more frequently to genes down-regulated in HepG2. **E** TTD binding to HepG2 chromatin is stronger around TSSs of genes that are most down-regulated in HepG2 compared to liver cells, and weaker around TSSs of genes that are up-regulated. The refTSS regions matching genes with ≥ 2-fold change in expression between HepG2 and liver tissue were arranged by increasing expression ratio (HepG2/Liver), divided in five bins of equal size, and the mean TTD signal of each was plotted. Central line is median, box borders are 25^th^ to 75^th^ percentile, and whiskers 5^th^ to 95^th^.

Using the ChIP-Enrich webserver (chip-enrich.med.umich.edu), a gene ontology biological process (GO:BP) analysis of the genes from cluster 1 of the TTD promoter binding analysis (Figure 5B) was conducted and revealed a strong connection to various metabolic processes, as well as response to xenobiotics (Figure 5C) (Supplemental File 2), both typical cell-type specific processes for hepatic cells ^54^. Then, we compared gene expression data from HepG2 and primary liver cells and identified genes with ≥ 2-fold change in expression and high expression levels in at least one of the two cell types (Supplemental File 3). We plotted the corresponding TSSs (refTSS) with their flanks, arranged by decreasing TTD signal, and noticed that TSSs with TTD-rich flanks corresponded more frequently to genes down-regulated in HepG2 (Figure 5D). Also, we arranged these TSSs according to change of expression, placed them in 5 bins of equal size and plotted the mean signal of TTD (Figure 5E), revealing that the TTD signal was stronger around TSSs of genes that are most down-regulated in HepG2 compared to liver tissue cells, and weaker around TSSs of genes that are up-regulated. Gene set enrichment analysis (GSEA) using Enrichr (maayanlab.cloud/Enrichr) ^55^ revealed that the genes robustly down-regulated between HepG2 and liver tissue are HepG2 and liver specific (Supplemental Figure 12A). These results show a correlation between UHRF1-TTD CIDOP signals on promoter flanks and reduced expression of a set of cell-type specific genes in HepG2 cells suggesting that these liver specific genes were downregulated by UHRF1 in the HepG2 cancer cells.

### UHRF1-TTD binds on enhancers of cell-type specific genes in HepG2

The enrichment of TTD peaks on ‘Enhancers’ in the chromatin segmentation analysis (Figure 5A) agrees with the overlap of TTD with H3K4me1. To examine the segmentation results closer, we merged the ∼ 189k ‘Enhancers’ of HepG2 cells and plotted a heatmap centered on these, with the marks relevant to our study (Figure 6A). As expected, all regions harbored a strong H3K4me1 signal, and a significant part of them also contains H3K9me2, showing a good correlation with the TTD signal (*r* 0.76). Direct comparison revealed that the majority of the TTD peaks (82%) are found on 30% of these HepG2-specific enhancers harboring H3K4me1 and H3K9me2 (Figure 6B). We also analyzed the TTD peaks on the ChIP-Enrich webserver (Supplemental Figure 12B) and determined that 65% of them are located on distal enhancers (10 to >100 kb to TSS) and 29% on upstream enhancers (1-10 kb to TSS), in agreement with our segmentation results. While the WGBS signal was lacking characteristic TTD-related pattern (Supplemental Figure 12C), a GO:BP analysis of the enhancer associated genes showed strong enrichment for processes related to metabolism and regulation of lipids followed by tissue/liver development (Figure 6C, Supplemental File 4). Among the enriched genes, we recognized liver specific markers (e.g. ALB, ALDOB, FGA) ^54^, as well as transcription factors (TFs) that define liver cell identity (e.g. FOXA1, HNF4A) ^56^ (Supplemental File 4). GSEA using Enrichr validated that the enriched assigned-genes are HepG2 and liver-tissue specific (Supplemental Figure 12D).

**Figure 6.**
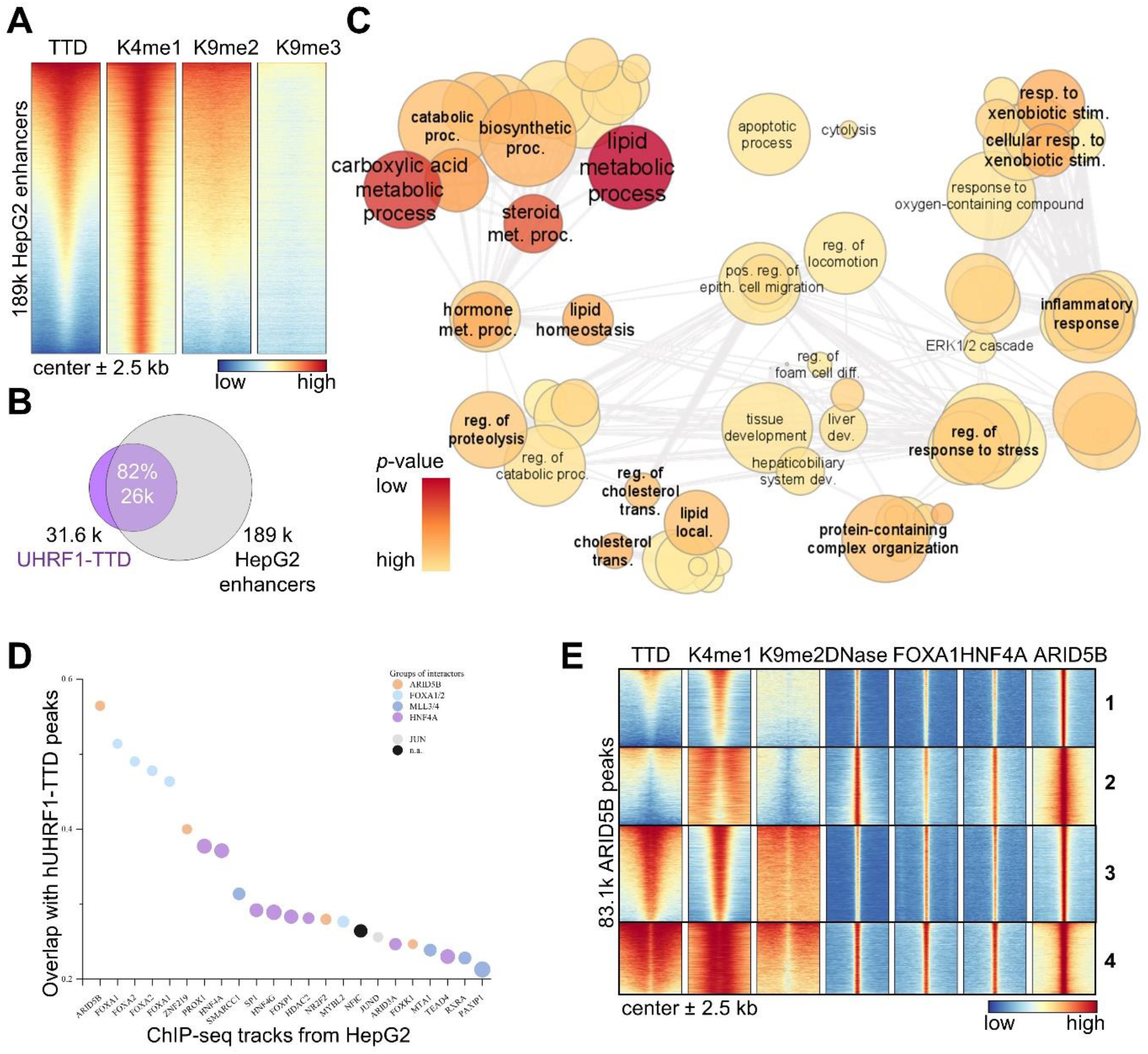
hUHRF1-TTD binds to enhancers of cell-type specific genes and flanks targets of cell-type specific TFs in HepG2. **A** Heatmap of ∼ 189k HepG2 enhancers identified by ChromHMM ± 2.5 kb flanks (egg2.wustl.edu/roadmap) ^51^, centered in the middle, and arranged by decreasing TTD signal. The TTD enrichment on HepG2 enhancers follows the H3K9me2 signal intensity. See also Supplemental Figure 12C. **B** UHRF1-TTD peaks have an 82% overlap with HepG2 enhancers, covering 30% of all HepG2 enhancers (ChromHMM). Diagram made using Venn-Diagram-Plotter (github.com/PNNL-Comp-Mass-Spec/Venn-Diagram-Plotter). See also Supplemental Figure 12B. **C** TTD peaks are enriched on enhancers of genes relating to cell-type specific processes for HepG2. TTD peaks were assigned to human enhancers by *ChIP-Enrich* (chip-enrich.med.umich.edu) ^85^, the enriched GO:BP genesets with FDR ≤ 0.05, hybrid p-value ≤ 0.05 were summarized in *Revigo* (revigo.irb.hr) ^86^ and visualized using *Cytoscape* (cytoscape.org) ^87^. Exemplary GO terms for each cluster are annotated. Semantic similarity is reflected in the clustering, color and font-size reflect *p*-value. Circle radius reflects the log of number of genes in GO term ID. dev -development; diff - differentiation; epith - epithelial; local - localization; met - metabolic; pos - positive; proc - processes; reg - regulation; res. – response; stim. – stimulus; trans - transport. See also Supplemental Figure 12D. **D** TTD peaks have strong correlation with targets of cell-type specific transcription factors (TFs). Overlap of TTD peaks with public ChIP-Seq peaks, individually curated to only include comparable HepG2 cells. Data shown here are ChIP-seq datasets with the highest overlap in TTD peaks in the public ChIP-Atlas database (chip-atlas.org) ^50^, as arranged by decreasing percentage of TTD peak overlap (counts). Each has log *p* < -10. Disk color reflects the known interactor/protein complex assigned to the specific protein. Disk size reflects the fold enrichment. **E** TTD flanks targets of cell-type specific TFs. Clustering revealed robust TTD binding surrounding ∼50% of the ARID5B peaks (clusters 3 and 4). Both show binding sites of DNA- binding cell-type specific TFs FOXA1 aka HNF3α ^89^, HNF4A ^89^, and ARID5B ^58^ and are flanked by H3K4me1-K9me2 and UHRF1-TTD. The center is nucleosome-free as seen by the DNase-seq signal ^53^. See also Supplemental Figure 12E.

### UHRF1-TTD and H3K4me1-K9me2 flank targets of cell-type specific TFs in HepG2

The TTD binding on enhancers regulating identity defining TFs (e.g. FOXA1, HNF4A), and the reported role of UHRF1 in regulating cell lineage specification during differentiation motivated further investigation in that direction. Analysis of peak overlap between TTD and TF ChIP-seq data from the ChIP-Atlas database for HepG2 cells ^50^, revealed a strong correlation of TTD to binding sites of cell-type specific TFs (Figure 6D, Supplemental File 5). Grouping of the most enriched TFs based on their known interaction (thebiogrid.org) ^57^ (Supplemental File 5) revealed the groups of ARID5B, FOXA1/2, MLL3/4, and HNF4A as most relevant TTD targets. Focusing on ARID5B, FOXA1 (aka HNF3α), and HNF4A, we verified that hUHRF1-TTD binds next to binding sites of cell-type specific TFs in browser views (Supplemental Figure 12E) and heatmaps (Figure 6E). The ChIP-seq data reveal the DNA-binding TFs are in the center of the TTD enriched regions, where nucleosomes are evicted, flanked by histone marks and TTD (e.g. cluster 4 of Figure 6E). The colocalization and interactions between these three TFs had been documented previously ^56, 58, 59^. Here, we document the presence of H3K4me1 and H3K9me2 at the flanks of these TF target regions, supporting the notion of a physiological role for this previously undescribed double mark and substantiating TTD binding to it. Moreover, the known interaction of FOXA1/2 with the MLL3/4 complex, that deposits H3K4me1, and HNF4A with G9a, that deposits H3K9me2, as well as with the MLL3/4 complex ^60, 61^ potentially leads to a double mark enhancement and may suggest a read/write mechanism of the H3K4me1-K9me2 double mark, since TF binding to regions containing H3K4me1-K9me2 can recruit writers of H3K4me1 and H3K9me2. In the cellular context, we expect that TTD-based targeting of UHRF1 to enhancers and promoters is subject to additional regulation, given the highly complex regulation of the interacting TFs. Taken together, our data clearly support a role for UHRF1 in differentiation and regulation of cell-type specific processes, mediated by TTD targeting. Our finding that H3K4me1-K9me2 and UHRF1-TTD flank the targets of cell-type specific TFs in HepG2 cells provides physiological context for this histone double mark, and a potential read/write mechanism via TFs and TTD, while providing an explanation for the previously reported role of UHRF1 in cellular differentiation ^34–38^.

### Murine UHRF1 genomic localization is correlated with H3K4me1

Next, we aimed to address the question, whether and to what extent the TTD data presented so far relate to the full-length UHRF1 protein. The only available full-length UHRF1 ChIP-seq data are from mouse embryonic stem cells (mESC). In a study that reported a genome-wide correlation of mUHRF1 with H3K4me3, as well as H3K9 methylation, and characterized mUHRF1 as a regulator of cell lineage specification during differentiation ^34^. After reanalysis of public E14 mESC tracks for H3K4me1 ^62^, H3K4me3 ^34^ and the mUHRF1 ChIP-seq data from Kim et al. (2018) ^62^, we concluded that the distribution of murine UHRF1 was much more similar to H3K4me1 than to H3K4me3 in browser views (Figure 7A) and the genome-wide correlation was better for H3K4me1 than for H3K4me3 (*r* 0.6 *vs*. 0.4) (Figure 7B, Supplemental Figure 13A). Evidently, the log2 plot of the H3K4me3 *vs*. mUHRF1 signal demonstrated a bimodal distribution (Supplemental Figure 13A), indicating that mainly the weaker peaks of H3K4me3 (which are expected to contain H3K4me1 as well) correlate with the UHRF1 signal in mESC, but strong H3K4me3 peaks showed no correlation with UHRF1.

**Figure 7.**
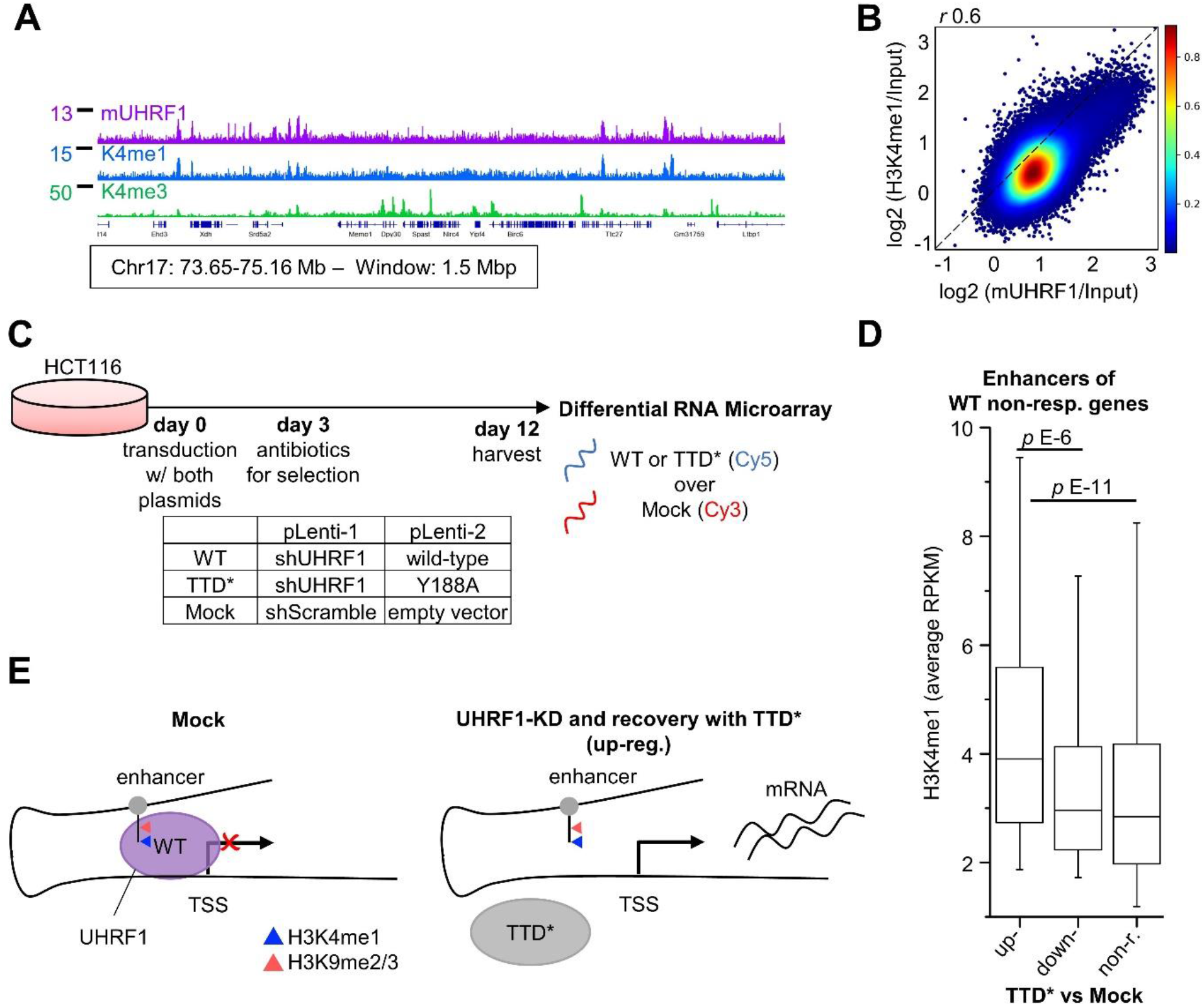
Full-length mUHRF1 genomic localization is correlated with H3K4me1 and hUHRF1-TTD down-regulates genes with H3K4me1 enriched enhancers. **A** Browser view of mUHRF1 ChIP-seq with H3K4me1 and H3K4me3 in E14 mESC demonstrates the similarity in signal and peak distribution. ChIP-seq datasets from Kim et al. (2018) ^34^ and Wu et al. (2016) ^62^. All tracks in RPKM, y-axes start from 0. Coordinates in mm10, gene annotation from RefSeq. Browser views of CIDOP-/ChIP-seq data were created with the Integrative Genomics Viewer (software.broadinstitute.org/software/igv/). **B** mUHRF1 ChIP-seq correlates with H3K4me1 in E14 mESC. Plot of the average ChIP signals in 2 kb bins genome-wide and Pearsońs correlation (*r*). Ratio calculated as log2(ChIP/Input). ChIP-seq datasets from Kim et al. (2018) ^34^ and Wu et al. (2016) ^62^. See also Supplemental Figure 13A. **C** Experimental strategy used by Kong et al. (2019) for the generation of differential expression data from modified HCT116 cells compared to mock to evaluate hUHRF1-TTD function ^64^. See also Supplemental Figure 13B. **D** TTD* up-regulated genes are enriched in H3K4me1-K9me2 on their FANTOM5 enhancers^65^. Shown are the differentially regulated genes (DRGs) from the wild-type (WT) over mock non-responsive genes, sorted according to their status in Y188A mutant over mock (TTD*). The mean H3K4me1 signal of each group was plotted from HCT116 ChIP-seq data ^66^. Central line is median, box borders are 25^th^ to 75^th^ percentile, and whiskers 5^th^ to 95^th^. *p* values are from one-way ANOVA with Bonferroni correction. non-r. – non-responding. See also Supplemental Figure 13C. **E** Schematic representation of TTD dependent regulation of gene expression in UHRF1 knock-down (KD) and Y188A (TTD*) rescued cells versus mock treated cells. Enrichment of the FANTOM5 enhancers in H3K4me1-K9me2 results in robust UHRF1 binding via TTD and down-regulation of the corresponding gene. Rescue with the H3-binding deficient mutant (TTD*) de-represses the gene.

As the direct interaction of UHRF1-TTD with the H3K4me1-K9me2/3 double mark discovered here provides the only known connection of UHRF1 to H3K4me1, we conclude that the correlation of the full-length murine UHRF1 ChIP-seq data with H3K4me1 strongly suggests that our observations with TTD are highly relevant for the chromatin interaction of full-length UHRF1 in cellular contexts. Although one specific splicing isoform of murine UHRF1 differs in chromatin binding from the human form ^63^, the observation of H3K4me1 dependent chromatin interaction for the murine UHRF1 and the human TTD strongly suggests that murine and human UHRF1 both bind to H3K4me1-K9me2/3.

### UHRF1-TTD down-regulates genes with H3K4me1-K9me2 enriched enhancers

Finally, we turned our attention to the potential physiological role of TTD binding to H3K4me1-K9me2/3 double marks. Previous work has shown the TTD dependent gene silencing by UHRF1 ^13^. To assess whether our finding of improved TTD binding to H3K4me1-K9me2/3 has implications in gene regulation, we turned to data published by Kong et al. (2019) ^64^. Facing the problem of toxicity of UHRF1 KO or KD, the authors generated HCT116 cells stably repressing endogenous UHRF1 with shRNA, which were simultaneously rescued with wild-type UHRF1 (WT) or a UHRF1-TTD mutant (TTD*) containing a Y188A mutation in the H3K9me2/3 binding pocket that disrupts H3-tail binding ^13^. As internal controls for the microarray analysis, mock-treated cells transduced with scrambled shRNA and empty vector were used (Figure 7C). Given the narrow dynamic range of the data, we used a modest threshold (|Fold Change over Mock| ≥ 1.5) to call differentially regulated genes (DRGs). The rescue with the TTD* mutant UHRF1 affected gene expression, as the WT UHRF1 rescued cells returned less DRGs than the TTD* UHRF1 rescued ones (Supplemental Figure 13B), but the discrepancy was small. To retain a robust gene-set, we first selected the 20911 genes that did not show an expression change after rescue with WT (when compared to mock treated cells) (Supplemental Figure 13B). This filters for genes where UHRF1 has no influence on expression or the UHRF1 rescue was fully functional. Among them, 115 were up-regulated after TTD* UHRF1 rescue (when compared to mock treated cells), indicating that the TTD domain is required for their silencing.

To investigate a potential connection between DRGs and H3K4me1 levels, we used the gene-specific enhancers from FANTOM5 ^65^, H3K4me1 and H3K9me2 data from HCT116 cells ^66, 67^, and plotted the mean signal in these regions. The 115 genes upregulated after reconstitution with TTD* UHRF1 were connected to enhancers that carry significantly more H3K4me1 (Figure 7D) and slightly less H3K9me2 than the non-responsive genes (Supplemental Figure 13C). The upregulation of these genes after reconstitution with TTD* UHRF1 indicates that they were originally repressed by UHRF1 in a TTD dependent manner (Figure 7E). The better correlation between the H3K4me1-K9me2 double mark and TTD-dependent gene silencing than with H3K9me2 alone agrees with all our previous data. Looking at the enhancers of all genes (not just the non-responders after reconstitution with WT UHRF1), we again found a higher H3K4me1 signal and marginal difference in H3K9me2 for the up-regulated DRGs (Supplemental Figure 13D). We conclude that genes with enhancers carrying H3K4me1-K9me2 are bound by UHRF1 via TTD and down-regulated (Figure 7E). UHRF1-knock down and rescue with the H3-binding deficient mutant (TTD* UHRF1) de-repressed these genes demonstrating the TTD dependent silencing of H3K4me1-K9me2 containing enhancers by UHRF1. This H3K4me1 dependent effect of human UHRF1 on gene regulation via TTD can only be explained in the context of the H3K4me1-K9me2/3 binding of TTD discovered in our work. It directly demonstrates that H3K4me1-K9me2/3 binding of TTD plays an important role in the cellular activities of full-length UHRF1.

Taken together, our findings indicate that the interaction of TTD with H3K4me1-K9me2 on enhancers is a driver for the UHRF1-mediated down-regulation of the corresponding genes. This directly associates the double mark H3K4me1-K9me2 and its interaction with TTD to a known physiological function of the full-length UHRF1. Our observation that DNA methylation is not detectably different in enhancer regions of HepG2 cells provides evidence that UHRF1 has a direct gene silencing role that is independent of its role in DNA methylation. This agrees with the findings of Kong et al. (2019), that Y188A has minimal effects on DNA methylation ^64^.

## Conclusions

Dissecting the roles and functions of the multidomain UHRF1 protein has not been an easy task in the two decades since its discovery ^68, 69^. UHRF1 is an essential chromatin factor needed for global maintenance of DNA methylation ^26–29^. It comprises different reading domains interacting with modified histone tails (TTD, PHD) and hemimethylated DNA (SRA), and has catalytic activity as ubiquitin ligase with its RING domain. Exploiting these activities, UHRF1 mediates several connections within the epigenome network and functions as a hub for recruitment of epigenetic effectors with a wide range of cellular functions ^26, 27^. However, even the individual building blocks of this essential master-regulator protein are insufficiently characterized so far. In this study, we discovered the combined H3K4me1-K9me2/3 readout of the UHRF1-TTD reading domain with both synthetic peptides and native nucleosomes. This is an interesting finding underscoring the results of previous reports that UHRF1 can interact with chromatin in different domain arrangements ^13, 40^, because in linked PHD-TTD structures, TTD binds H3K9me3 and PHD binds H3R2 while the linker peptide between both domains was observed to block the H3K4me1 binding pocket of TTD identified here ^20, 22^. Indeed, changes in the domain arrangement of full length UHRF1 upon binding to the LIG1 peptide have been directly observed in SAXS experiments ^18^ and phosphorylation of a linker residue was implicated in altered domain arrangements of UHRF1 as well ^20^. Moreover, the H3K4me1 binding pocket on TTD has been shown to mediate the preferable interaction of full length UHRF1 with R121 in LIG1 ^18, 19^ which has well documented physiological roles in cells, indicating that this binding site is available in full-length UHRF1 in cells. *In vitro* data strongly support this, as the TTD cleft occupancy by the linker was reported to be approximately 50%^70^.

Using TTD mutants, we demonstrated that binding to the H3K4me1-K9me3 peptide makes use of a novel Kme1 recognition process that has two contributing principles. Firstly, it is based on the reduced H-bonding potential of Kme1 when compared to Kme0 and secondly on the vdW interactions of the Kme1 methyl group with the methylene groups of an arginine residue in the TTD. Strikingly, the E153A TTD mutant showed an 10,000-fold preference for the double modified peptides, suggesting that conformational changes moving E153 out of the peptide binding cleft could lead to similar preferences of WT TTD, which could explain the strong H3K4me1-K9me2/3 preferences of TTD observed in our chromatin binding experiments and in literature UHRF1 ChIP-seq data. This finding suggests that conformational changes of UHRF1 mediated by PTMs or binding of other proteins could regulate its binding to double H3K4me1-K9me2/3 *vs.* isolated H3K9m2/3, an interesting hypothesis that needs to be further investigated.

Having shown preferential TTD binding to HepG2 native nucleosomes carrying H3K4me1-K9me2/3 double modifications, we also show correlation of available full-length mUHRF1 ChIP-seq profiles to H3K4me1, indicating that our observations are relevant for full-length UHRF1 chromatin binding in its biological context. On the genome-wide scale, we show that the H3K4me1-K9me2 double mark is a frequent modification suggesting that it has a specific role. UHRF1-TTD binds to the flanks of gene promoters for cell-type specific processes and is enriched on the flanks of genes with strongly down-regulated expression suggesting a H3K4me1-K9me2/3 dependent repressive role of UHRF1. This agrees with a previous report that UHRF1 binds gene promoters and mediates silencing of the associated genes in cancer cells ^27^. Moreover, we observed that UHRF1-TTD is enriched in enhancers with a strong connection to genes of cell-type specific processes and cell-type specific TFs in hepatic cells, in line with reports of UHRF1 involvement in cell-type specific gene regulation during lineage specification ^34–38^. Conversely, the enhancers of genes down-regulated by full-length hUHRF1 in a TTD dependent manner in HCT116 cells demonstrate an enrichment in H3K4me1-K9me2, indicating that UHRF1 contributes to silencing of H3K4me1-K9me2 marked enhancers. These data again demonstrate the direct relevance of H3K4me1-K9me2 binding by UHRF1-TTD and its physiological function in a cellular context. However, future studies need to further address the role of H3K4me1 binding by UHRF1 *in vivo*, for instance in the UHRF1 mediated repression of tumor suppressor genes such as p16INK4A ^71, 72^.

Our novel finding of the preferential binding of UHRF1-TTD to H3K4me1-K9me2/3 can also directly explain previous observations of Skvortsova et al. (2019) that the presence of H3K4me1 at CpG island borders predisposes these regions for gain in DNA methylation ^73^. In light of our findings, these data could be explained by increased recruitment of UHRF1 due to the presence of H3K4me1-K9me2/3. Our new data will assist future studies on the functions and effects of UHRF1 to comprehensively describe the multifaceted biological functions of this important chromatin factor, which represents an important node in the epigenome network ^26, 27, 74^ and a known oncogene in liver and other carcinomas ^74, 75^.

## Materials and Methods

### GST-recombinant proteins

The Tandem Tudor domain of hUHRF1 (UNIPROT Q96T88) on previously defined borders (residues 126-280) ^14^ was N-terminally fused to GST, under control of a *lac* promoter. Mutations were introduced using an updated rolling circle protocol and validated by Sanger sequencing. The oligonucleotides used for mutagenesis are listed in Supplemental Table 1. Proteins were overexpressed in *E. coli* by induction with 1 mM isopropyl-β-D-thiogalactopyronoside (IPTG) when OD_600_ reached 0.6 to 0.8, and the culture was continued overnight at 20 °C. The cells were harvested by centrifugation for 30 min, at 4 °C and 3,781 *g*. For purification, each pellet was resuspended in 30 ml sonication buffer (500 mM KCl, 20 mM HEPES pH 7.5, 10% v/v glycerol, 0.2 mM dithiothreitol (DTT)) and sonicated for cell lysis (Q120 Sonicator, Active Motif). After centrifugation for 1 h at 4 °C and 45,850 *g*, the soluble protein was purified using Glutathione Agarose 4B gravitational columns and dialysed against dialysis buffer (200 mM KCl, 20 mM HEPES, 10% v/v glycerol, pH 7.5, 0.2 mM DTT). Protein aliquots were flash-frozen and stored at -80 °C. Protein concentrations were determined spectroscopically by A_280_ and purity was verified on SDS-PAGE.

### CelluSpots array binding experiments

MODified™ Histone Peptide Arrays (Active Motif) were processed according to previously published protocols ^39, 43, 45^. Briefly, the array was blocked in 5% w/v skim milk in TBS with 0.1 % v/v Tween20, then incubated with 500 nM GST-hUHRF1-TTD in interaction buffer (100 mM KCl, 20 mM HEPES pH 7.5, 10 % v/v glycerol) at room temperature. Anti-GST (GE Healthcare, #27457701V) and anti-goat-HRP (Sigma Aldrich, #A4174) antibodies were used for visualization. Antibody specificity validation was performed using CelluSpots arrays and the antibodies diluted in 1% w/v skim milk in TBST, as reported in Supplemental Table 3. Processing was as described above, with appropriate secondary antibodies. Detailed information on the primary antibodies and lot specific validation data are given in Supplemental Figure 4 and Supplemental Table 3.

### Equilibrium peptide binding titrations

Equilibrium peptide binding experiments were performed on a Jasco FP-8300 spectrofluorometer with an automatic polarizer FDP-837. The FITC labelled peptides for fluorescence anisotropy (FA) titrations were purchased from commercial vendors (Supplemental Table 2). Peptides were purified by RP-HPLC and had a final purity of ≥ 90%. Acquisitions were performed at 23 °C with excitation at 493 nm and emission measured at 520 nm, slit width set to 5 nm for both, and multiple accumulations. FA buffer consisted of 100 mM KCl, 20 mM HEPES pH 7.5, 10% v/v glycerol. The initial concentration of the fluorescent peptide was 100 nM. The protein was added stepwise to the cuvette. Titrations were replicated at least two times in independent experiments. Data processing was performed with Microsoft Excel. To determine K_D_ values, the data were fitted to a simple binary equilibrium binding model (Equation 1):

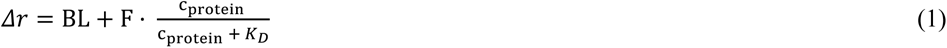

where Δr is the anisotropy signal, BL is baseline, F is signal factor, K_D_ is the equilibrium binding constant, and c_protein_ refers to total concentration of protein.

For K_D_ values below 100 nM, the data were fitted to an expanded binary equilibrium binding model (Equation 2):

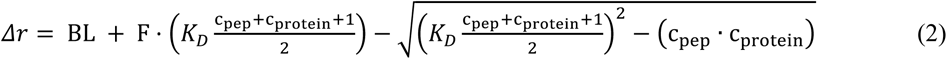

where Δr is the anisotropy signal, BL is baseline, F is signal factor, K_D_ is the equilibrium binding constant, c_protein_ refers to the total concentration of protein, and c_pep_ refers to the total concentration of peptide.

### CIDOP and ChIP

HepG2 cells were acquired from DSMZ - German Collection of Microorganisms and Cell Cultures (No: ACC 180) and grown in RPMI 1640 medium (Gibco) supplemented with 10% FBS, 100 U/ml penicillin and 100 mg/ml streptomycin at 37 °C under humidified air with 5% CO_2_. Cells were harvested at 300 *g* (5 min, 4 °C) and the pellets were washed once with 1 ml PBS, flash-frozen and stored at -80 °C. For CIDOP-Western blot, biological triplicates or better were generated using separately cultured HepG2 cells. For ChIP-seq, and CIDOP-seq, biological duplicates were generated. Mononucleosome generation and histone precipitation were performed with a modular protocol, all parameters optimized for each enrichment reagent (Supplemental Table 4). Briefly, the cells were resuspended in lysis buffer (10 mM Tris-HCl pH 7.4, 2 mM MgCl_2_, 0.5 mM PMSF, 1 mM DTT, 0.6% v/v Igepal CA-360, EDTA-free protease inhibitor cocktail tablet), digested with ∼ 135 units of MNase (NEB, M0247) per 1 million cells at 37 °C, 150 rpm for 12.5 min in one tube, diluted in interaction buffer (20 mM Tris-HCl pH 8.0, 150 mM NaCl, 1 mM PMSF, 0.1% v/v Triton X-100, 50% v/v glycerol, EDTA-free protease inhibitor cocktail tablet) centrifuged, and the supernatant containing mononucleosomes collected, flash-frozen and stored in -80 °C. HiMIDs or α-H3K9me2 (ab1220) were first incubated with appropriate magnetic beads (GST-Pierce magnetic or DynabeadsG 10004D) for 2 h, and then with precleared chromatin for overnight binding. Beads were washed three times with PB200 (50 mM Tris-HCl pH 8.0, 200 mM NaCl, 2 mM DTT, 0.5% v/v Igepal CA-360), followed by two rinse steps (10 mM Tris-HCl pH 8.0 and optionally 150 mM LiCl). Samples for western blot were then heated to 95 °C in loading buffer (160 mM Tris-HCl pH 6.8, 2% w/v SDS, 40% v/v glycerol). The wet transfer protocol was optimized for H3 histones, using MeOH-free Towbin buffer (25 mM Tris, 192mM glycine, 0.02% SDS w/v, and 20% EtOH v/v). Samples for qPCR/NGS analysis were eluted (50 mM Tris-HCl pH 8.0, 50 mM NaCl, 5mM EDTA, 1% w/v SDS), digested with 2.5 units of Proteinase K (NEB, P8107) at 55 °C, 900 rpm for 90 min, and purified with the ChIP DNA Purification Kit (Active Motif). The qPCR assays were performed on a CFX96 qPCR system (Bio-Rad) using ORASEE qPCR reagent (highQu). The oligonucleotides used for qPCR assays are listed in Supplemental Table 5. NGS libraries were prepared with 10 ng DNA using the NEBNext® Ultra™ II DNA Library Prep Kit according to the manufacturer’s instructions and sequenced on an Illumina NovaSeq 6000 with 150 bp paired-end reads for a minimum of 10 million reads.

### CIDOP-seq and ChIP-seq data analysis

Data analysis was performed on a *Galaxy* server (usegalaxy.org) ^76^. Publicly available ChIP data were obtained as raw reads from SRA (ncbi.nlm.nih.gov/sra), accession codes are given in Supplemental Table 6. The *deeptools2* ^77^ and *bedtools* ^78^ suites, as well as *ChAsE* ^79^ and *Integrative Genomics Viewer* (IGV) (software.broadinstitute.org/software/igv/) ^80^ software were used for downstream data processing and visualisation. Browser views of CIDOP-seq and ChIP-seq data were created with the Integrative Genomics Viewer (software.broadinstitute.org/software/igv/). Adapters were clipped and low-quality reads removed with *Trimmomatic* (v0.38) using default settings, and quality controlled with *FastQC* (v0.72) ^81^. The high-quality, clean reads were mapped to hg38 or mm10 using *HISAT2* (v2.2.1)^82^. Using *bamcoverage* (v3.3.2), the mapped reads were quantified in 10 bp bins using Reads Per Kilobase of transcript, per Million mapped reads (RPKM), omitting blacklisted regions (hg38- or mm10-blacklist.v2) ^46^. Biological replicates were pooled using *bigwigcompare* (v3.3.2) and the mean RPKM signal determined. Pearson correlation factors were calculated with *deepTools2*, using 2 kb bins for genome wide comparisons of UHRF1-TTD CIDOP-seq pooled data to the individual replicates and the H3 PTMs. To compare pooled data to the individual replicates for the very broad H3K9me2 mark, 10 kb bins were used.

### Splitting of the genome in deciles

The hg38 genome excluding blacklisted regions was separated in 1 kb bins using *MakeWindowsBed* (v2.30), and the average signal in each bin was computed using *multibigwigSummary* (v3.3.2). The regions were ranked by descending H3K9me2 signal and split into 10 groups, each with an equal number of regions (286110), representing the 10 deciles. For all box plots, the central lines show the median, box borders are 25^th^ to 75^th^ percentile, and whiskers 5^th^ to 95^th^. Pearson correlation scores were calculated with *deepTools2*, using the average values within the 1kb bins.

### CIDOP peak calling and fragmentation

Broad peaks for TTD were called with *MACS2* (v2.1.1) using cut-off 30, cut-off-link 13, d 150, t 89. Blacklisted regions were removed and the peaks were manually curated for artefacts and large, false positives. To fragment the UHRF1-TTD peaks, the hg38 was split into 150 bp bins using *MakeWindowsBed* and those with a ≥ 50% overlap with TTD peaks were selected using *MapBed* (v2.30). *k*-means clustering was performed using *ChAsE*.

### Heatmaps and k-means clustering

Bed files used for heatmaps were arranged by descending TTD intensity, and *k*-means clustering was performed using *ChAsE*. Heatmaps were generated using *deepTools2*. For box plots, average signals in each region were computed using *multibigwigSummary*. WGBS signals are depicted with the same color-range (min-max) in both heatmaps. Pearson correlation factors were calculated with *deepTools2*, using the average values within the peaks for Figure 4D and E. For Figure 6A, due to the variability of enhancer size, the average values within the 5 kb window were used.

### Histone PTM and TF peak overlap analysis

To investigate the overlap of TTD peaks with histone PTM or TF ChIP-seq peaks, the ChIP-atlas database (chip-atlas.org) enrichment analysis tool was used ^50^. The search was restricted to liver cells and the control was a 10x genome-wide, random permutation of the peak file. The results were further restricted to data from HepG2 cells, and ≥ 1.0-fold enrichment (Supplemental Files 1 and 5). The results contained data from experiments with wild-type cells, but also knock-downs/-outs, transfected and treated cells, as well as non-typical ChIP techniques (low input etc). Data with ≥ 4.0-fold enrichment were individually curated to originate only from wild-type cells ChIP experiments and study abundant histone PTMs. Similarly, the TF ChIP-seq results were individually curated to only include data from comparable HepG2 cells.

### Murine ChIP analyses

To generate similar plots as Kim et al. 2018, *bigwigcompare* was used to report the log2 ChIP over input signal from the mm10 mapped, RPKM normalized bigwig files. The average signal in 2 kb bins was computed using *multibigwigSummary* excluding blacklisted regions. Pearson correlation factors were calculated with *deepTools2*, using 2 kb bins for the genome-wide comparisons. Scatter-plots were generated using *MatPlotLib*.

### Chromatin segmentation analysis

The 18-state ChromHMM data for HepG2 cells were obtained from egg2.wustl.edu/roadmap ^51^. A control file with regions of equal number and equal length to the TTD peak file was created using *shuffle bed*. Using *Annotate bed*, overlaps of the ChromHMM regions with TTD peaks and the control were counted and the ratio of observed over expected (TTD/control) calculated for each of the 18 states.

### Promoters and expression levels

For the TSS regions, the refTSS (v3.1_hg38) regions were used ^52^. WGBS data were downloaded as a pre-processed bigwig file from ENCODE ^53^ (Supplemental Table 6). Analysis of the expression levels in HepG2 and liver tissue cells was done using pTPM (protein-transcripts per million) data from The Human Protein Atlas version 21.1 (proteinatlas.org) ^83, 84^. The genes were selected for ≥ 2-fold change in expression (HepG2/Liver) and a high expression level (≥ 1000 pTPM in one of the two cell types), to avoid false positives and small effects (Supplemental File 3). For the box plot, the regions were arranged by increasing expression ratio (HepG2/Liver), the average TTD signal within the 5 kb window was computed using *multibigwigSummary*, and the regions distributed in 5 bins of equal size. For all box plots, the central line is median, box borders are 25^th^ to 75^th^ percentile, and whiskers 5^th^ to 95^th^.

### Enhancer heatmap

The regions from all the “Enhancers” states were selected to create the “HepG2 enhancers” bed file from the HepG2-specific 18-state ChromHMM reference data (egg2.wustl.edu/roadmap)^51^. Pearson correlation scores were calculated with *deepTools2*, using the average values within the 5 kb window, due to the variability of enhancer size.

### Enrichment analyses

For the TSSs of cluster 1 (Figure 5B), we performed TSS-to-gene assignment and GO:BP analysis of these genes, using the ChIP-Enrich method on the *ChIP-Enrich* webserver (chip-enrich.med.umich.edu) ^85^ with the settings nearest TSS and adjust for mappability – true. The resulting GO assignments were filtered for FDR ≤ 0.05, *p*-value ≤ 0.05, status enriched (Supplemental File 2). The geneset ID with the corresponding *p*-value were summarized by in *Revigo* (revigo.irb.hr) ^86^ for a small list of GO terms. The resulting network was visualized using *Cytoscape* (cytoscape.org) ^87^. For enhancer-to-gene assignment and GO:BP analysis, we used the hybrid method of *ChIP-Enrich* ^88^ with the settings nearest gene and adjust for mappability – true. The resulting GO assignments were filtered for FDR ≤ 0.05, hybrid *p*-value ≤ 0.05, status enriched (Supplemental File 4). These were summarized to a small list by *Revigo*, and visualized using *Cytoscape*.

### TF and DNase data

ARID5B ^58^, FOXA1 ^89^, HNF4A ^89^ ChIP-seq data from HepG2 cells were retrieved from SRA and processed as described in “CIDOP-seq and ChIP-seq data analysis”. ARID5B peaks for hg38 were retrieved from chip-atlas.org, and used without additional processing. DNase data for HepG2 cells ^53^ were retrieved as bigwig files from ENCODE and merged for average signal as described in “CIDOP-seq and ChIP-seq data analysis”, and used without additional processing.

### Gene Expression Microarrays

Pre-processed differential expression data from Agilent human gene expression microarrays (Agilent4x44K v2 G4845A 026652) were downloaded from GSE118971 and processed as described previously ^64^. In brief, the pre-processed Lowess normalized log_2_(Fold Change) ratio was matched to the corresponding gene name and the median calculated from the ≥ 1 probes within each gene. To retain an adequate number of genes we used a moderate cut-off to call differentially regulated genes (median |FC| ≥ 1.5) compared to mock cells.

### FANTOM5 enhancers

FANTOM5 enhancers assigned to genes were retrieved from the FANTOM5/PrESSTo database (enhancer.binf.ku.dk/presets/enhancer_tss_associations.bed) ^65^, lifted over to hg38, and sanitized. H3K4me1 and H3K9me2 ChIP-seq data from HCT116 cells ^66, 67^ were retrieved from SRA and processed as described in “CIDOP-seq and ChIP-seq data analysis”. For the box plots, the average H3K4me1 signal within each enhancer region was computed using *multibigwigSummary*, and plotted. For all box plots, the central line is median, box borders are 25^th^ to 75^th^ percentile, and whiskers 5^th^ to 95^th^.

### Statistics

Standard deviations were calculated using the STDEV.P command in Excel. Confidence intervals were calculated using the CONFIDENCE.NORM command at the 0.05 significance level. For equality of variances on two experimental conditions we used F-test and for *p*-value calculation we used the unpaired, two-tailed Student’s t-test (H_0_: difference of means = 0, α = 0.05) with or without assumption of equal variances, as appropriate. Significance levels were assigned as follows: n.s. *p* > 0.05, \**p* ≤ 0.05, \*\**p* ≤ 0.01. For multiple comparisons, *p*-values were calculated using one-way ANOVA with Bonferroni correction.

### Data availability

The UHRF1-TTD CIDOP-seq and H3K9me2 ChIP-seq data are available at GEO under the accession number GSE213741. Anonymous reviewer access to this entry is available at: https://www.ncbi.nlm.nih.gov/geo/query/acc.cgi?acc=GSE213741

Using the access token: kruzogigvjghtcj

## Supplementary material description

### Supplemental Information

Supplemental Figures 1-13, Supplemental Tables 1-6 and Supplemental references

### Supplemental data provided as Excel files

Supplemental File 1. UHRF1-TTD and H3 PTMs overlaps in ChIP-Atlas database.

Supplemental File 2. ChIP-Enrich results for UHRF1-TTD enriched refTSS-cluster 1.

Supplemental File 3. TSSs of genes with ≥ 2-fold change in expression (HepG2 *vs.* Liver).

Supplemental File 4. UHRF1-TTD peaks analyzed by ChIP-Enrich-hybrid.

Supplemental File 5. UHRF1-TTD and TF overlaps in ChIP-Atlas database.

## Acknowledgments

MC thanks Julian Broche for training in NGS data analysis and insightful discussions. MC acknowledges the efforts of all involved in creating, maintaining and financing public data, databases and processing tools.

## Author contributions

MC and AJ devised the study. MC conducted all experiments and bioinformatic analyses in this study. RM and GK cloned the expression construct of TTD and made the initial observation. PB contributed to the bioinformatic work. MC, PB and AJ were involved in data analysis and interpretation. PB and AJ supervised the study. MC and AJ prepared the figures and first draft of the manuscript. All authors approved the final version of the manuscript.

