## Supplemental Information for "Refined read-out: The hUHRF1 Tandem-Tudor domain prefers binding to histone H3 tails containing K4me1 in the context of H3K9me2/3"

#### Supplemental Figures

Supplemental Figure 1. Additional data related to Figure 1.

Supplemental Figure 2. Influence of arginine methylation on binding of H3K9me2/3 peptides by TTD.

Supplemental Figure 3. Controls and exemplary primary data of the hUHRF1-TTD peptide binding experiments shown in Figure 2 and Table 1.

Supplemental Figure 4. Antibody validation data related to Figure 3B and Supplemental Figure 5A.

Supplemental Figure 5. Additional data related to Figure 3B.

Supplemental Figure 6. Validation of CIDOP-seq and ChIP-seq data shown in Figure 3C by CIDOP-qPCR.

Supplemental Figure 7. Additional browser views of the CIDOP-seq and ChIP-seq data shown in Figure 3C demonstrating reproducibility of experimental repeats.

Supplemental Figure 8. Additional browser views of the CIDOP-seq data shown in Figure 3C demonstrating TTD CIDOP signal enrichment in regions with both H3K4me1 and H3K9me2/3.

Supplemental Figure 9. Additional browser views of the ChIP-seq data shown in Figure 3E demonstrating the broad genomic localisation of H3K9me2 and H3K9me3.

Supplemental Figure 10. Additional data related to Figure 4.

Supplemental Figure 11. Additional data related to TTD binding to H3K4me1-K9me3 shown in Figure 4.

Supplemental Figure 12. Additional data related to promoter and enhancer binding in HepG2 cells.

Supplemental Figure 13. Additional data related to full-length mUHRF1 colocalization with H3K4me1 and gene regulation by hUHRF1-TTD in HCT116 cells shown in Figure 7.

#### Supplemental Tables

Supplemental Table 1. Oligonucleotides used for mutagenesis in this study.

Supplemental Table 2. Fluorescent histone H3 peptides used in this study.

Supplemental Table 3 Antibodies used for ChIP and/or western blots, with conditions of use.

Supplemental Table 4. Reagents and conditions used for CIDOP and ChIP from HepG2 mononucleosomes.

Supplemental Table 5. Oligonucleotides used for qPCR assays in this study.

Supplemental Table 6. NGS public datasets used in this study.

**Supplemental data provided as Excel files**

Supplemental File 1. UHRF1-TTD and H3 PTMs overlaps in ChIP-Atlas database.

Supplemental File 2. ChIP-Enrich results for UHRF1-TTD enriched refTSS-cluster 1.

Supplemental File 3. TSSs of genes with ≥ 2-fold change in expression (HepG2 *vs.* Liver).

Supplemental File 4. UHRF1-TTD peaks analyzed by ChIP-Enrich-hybrid.

Supplemental File 5. UHRF1-TTD and TF overlaps in ChIP-Atlas database.

**Supplemental references**

### Supplemental Figures

#### Supplemental Figure 1. Additional data related to Figure 1.

**A** Bar diagram of the average signal intensities for the ten best-bound peptides arranged by decreasing average signal, from two independent replicates of CelluSpots with UHRF1-TTD each providing two independent assays (Figures 1B and C). Bars represent the mean and error bars the standard deviation, n=4.

**B** Overlay of H3-K9me3 and LIG1-K126me3 peptides bound to UHRF1-TTD (PDB: 2L3R and 5YY9).

**C** Overlay of UHRF1-TTD acidic pocket residues in the H3-K9me3 and LIG1-K126me3 complexes. Panels (B) and (C) were generated with Chimera v1.14 (rbvi.ucsf.edu/chimera).

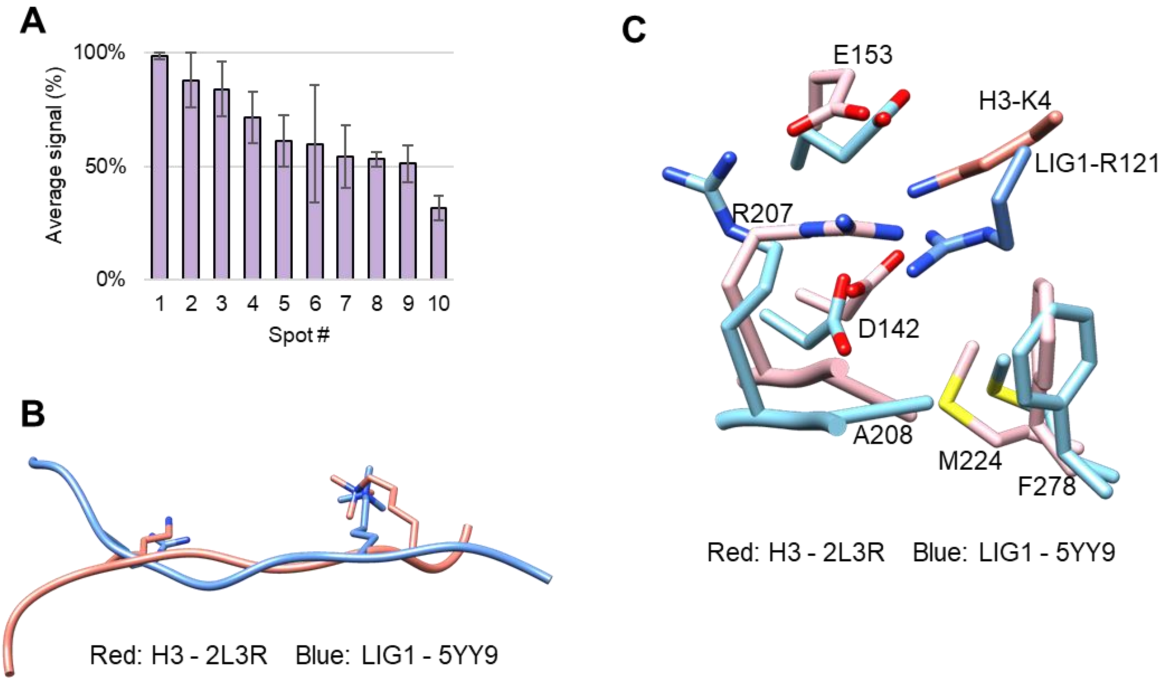

**Supplemental Figure 2. Influence of arginine methylation on binding of H3K9me2/3 peptides by TTD.**

**A** Structure of the UHRF1 TTD-LIG1 peptide complex ([1]. PDB: 5YY9). To investigate the potential influence of R-methylation on TTD peptide binding, we examined crystallographic data. TTD is shown in tan as surface. The peptide is shown in orange. R121 corresponding to H3K4 and K126me3 corresponding to H3K9me3 are shown in blue and pink. They approach the protein in two distinct deep binding pockets. P119 corresponding to H3R2 and R125 corresponding to H3R8 are shown in yellow. These residues point into the solvent (P119) or are placed on the surface of the domain (R125) without an apparent opportunity for preferential Rme2 binding.

**B** As shown in panel (A), the H3R8 equivalent residue makes a surface contact to the protein, hence the effect of its methylation has been studied in more detail. TTD binding to H3 peptides with K9me2 on Celluspots peptide arrays was inhibited by both structural isomers of R8me2 (symmetric and asymmetric). Unfortunately, no H3R2me-K9me2/3 double mark peptides are available on CelluSpots arrays.

**C** hUHRF1-TTD binding to the H3R8me2s-K9me3 peptide was studied in equilibrium peptide binding titrations analysed by the fluorescence anisotropy (FA) change and shown to have a $K_D$ of 2400 nM, while the $K_D$ of binding to H3K9me3 is 680 nM. This result inidcates that H3-tail binding is inhibited in presence of R8me2s.

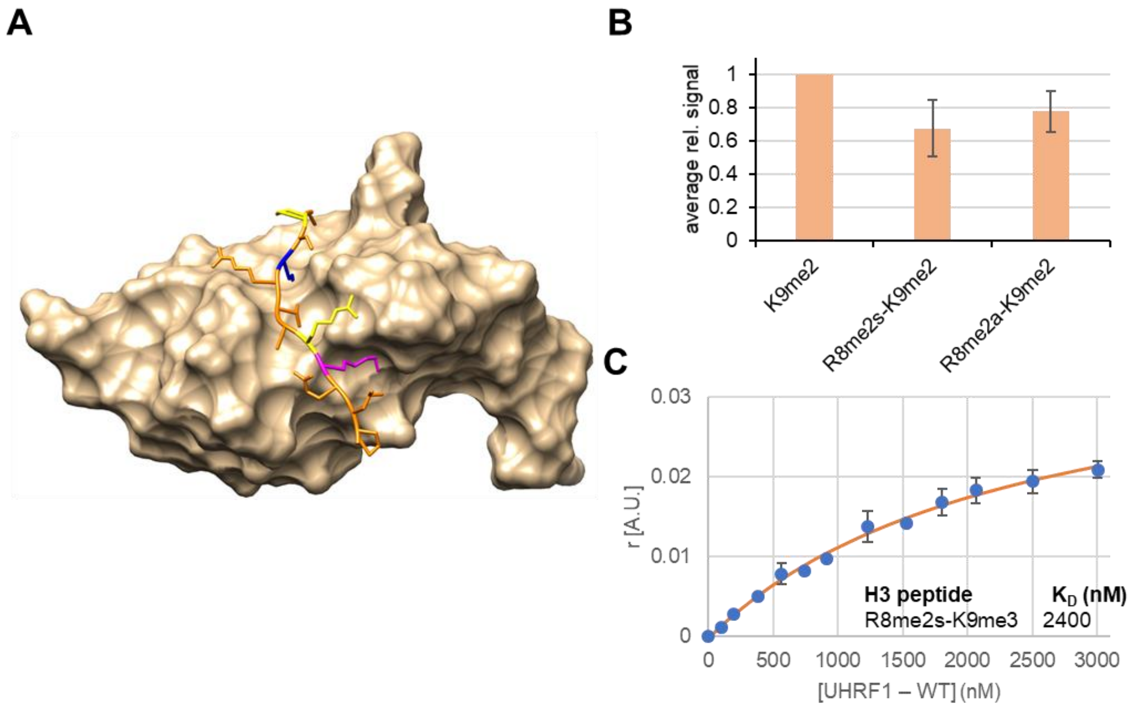

**Supplemental Figure 3. Controls and exemplary primary data of the hUHRF1-TTD peptide binding experiments shown in Figure 2 and Table 1.**

**A** TTD wild-type (WT) binds H3K4me1-K9me2/3 peptides more strongly than H3K9me2/3 alone in equilibrium peptide binding titrations analysed by the fluorescence anisotropy (FA) change.

**B** Control data showing that TTD WT binds weakly to the H3K4me1 peptide.

**C** Representative data showing peptide binding of TTD M224A.

**D** Representative data showing peptide binding of TTD D142A.

**E** Representative data showing peptide binding of TTD D142E.

Data points are average fraction bound (Θ) of $n \geq 2$ independent experiments, error bars are 0.95 confidence intervals (CI). $K_D$ are the mean of $n \geq 2$ independent fits, errors are 0.95 confidence intervals (CI).

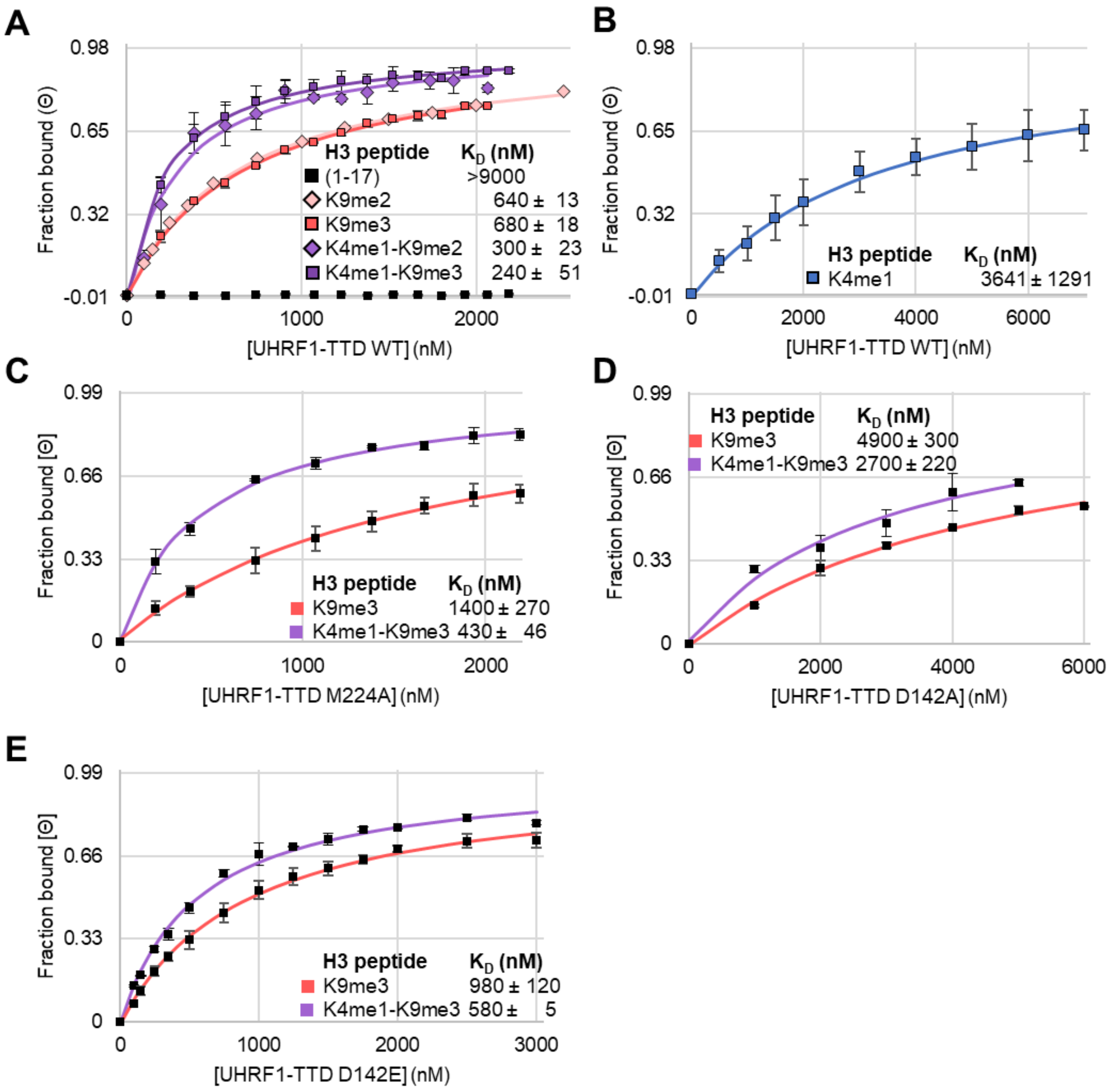

**Supplemental Figure 4. Antibody validation data related to Figure 3B and Supplemental Figure 5A.**

Lot-specific validation of antibody specificity using CelluSpots arrays, performed in the same time frame as the corresponding western blot experiments. All used antibodies are specific for the correct H3 PTM.

**A** Antibody validations for the CIDOP experiments shown in Supplemental Figure 5A. The α-K4me1 was tested under ChIP conditions and the α-K9me2 under western blot conditions to validate the antibodies' specificity in sequential ChIP-western blot. Catalogue and lot numbers are on the top left of the corresponding array. The target epitope is indicated in the box above the array and peptides with the epitope are marked with a circle of the same colour.

**B** Antibody validations for the CIDOP experiments shown in Figure 3B. Experiments were conducted under western blot conditions (Supplemental Table 3).

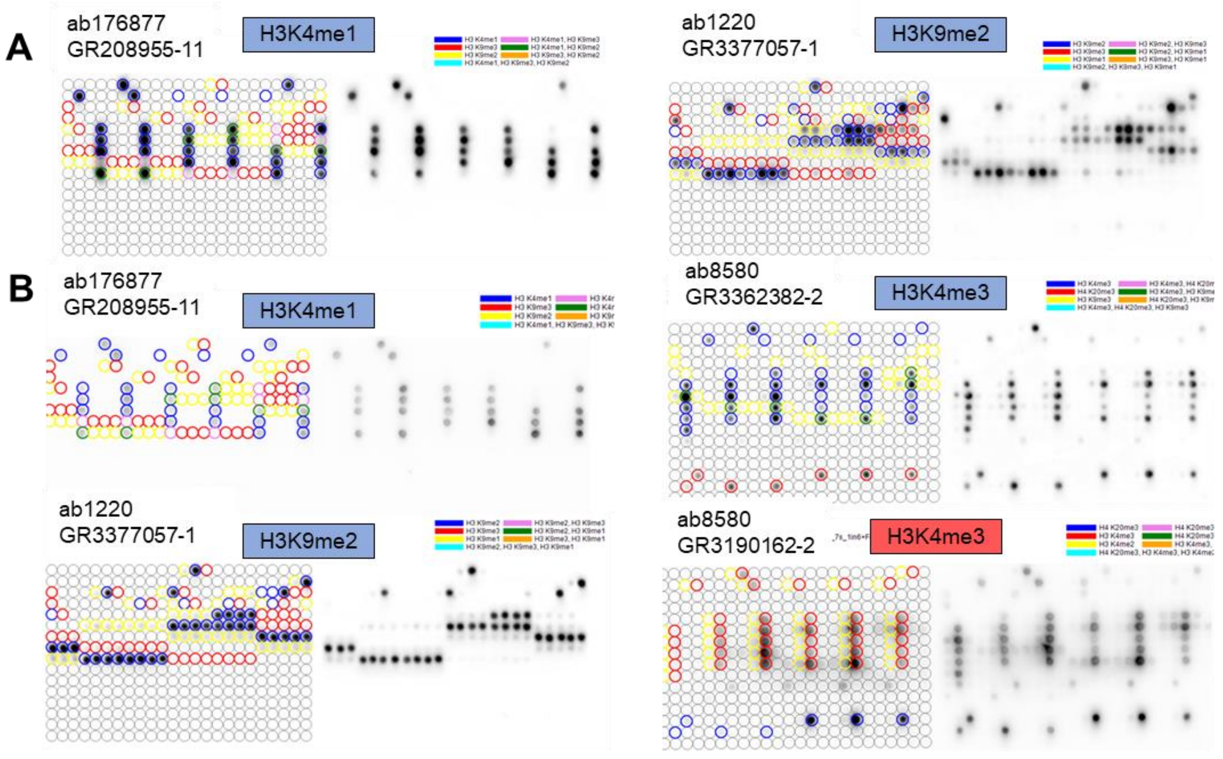

**Supplemental Figure 5. Additional data related to Figure 3B.**

**A** Detection of the occurrence of the H3K4me1-K9me2 double mark by H3K4me1 ChIP followed by H3K9me2 western blot. Primary data of western blots from 5 biological replicates of the α-K4me1 ChIP experiments.

**B** Quantification of western blots shown in panel A from $n = 5$ biological replicates of α-K4me1 ChIP experiments. The bars show the mean and error bars 0.95 confidence intervals.

**C** Primary data of western blots from 3 additional biological replicates of the CIDOP experiments shown in Figure 3B.

**D** Quantification of western blots shown in panel C from $n \geq 3$ biological replicates of CIDOP experiments. The bars show the mean and error bars 0.95 confidence intervals (CI). Significance levels were assigned as follows: n.s. $p > 0.05$, $*p \leq 0.05$, $**p \leq 0.01$.

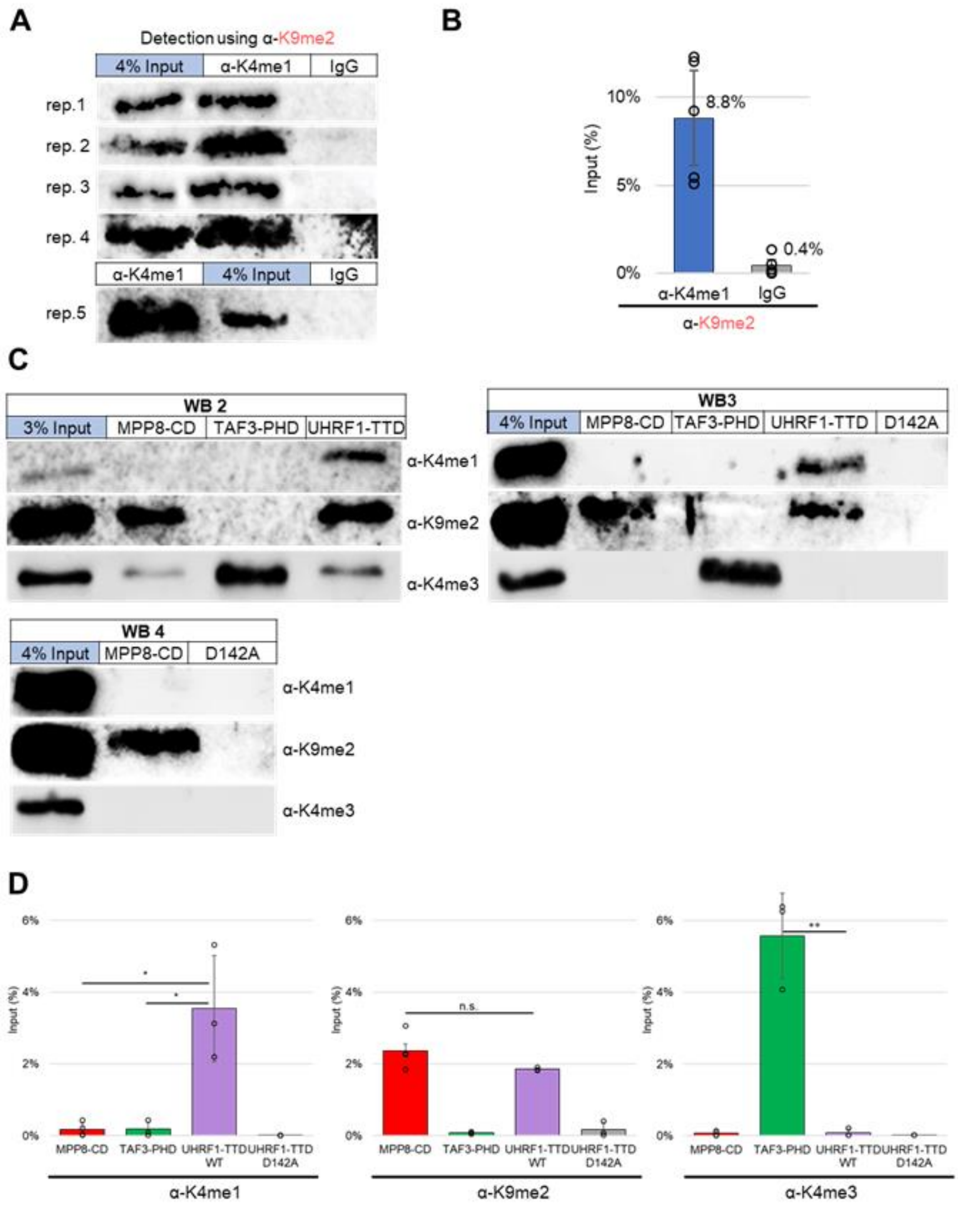

**Supplemental Figure 6. Validation of CIDOP-seq data shown in Figure 3C by CIDOP-qPCR.**

**A** TTD CIDOP-qPCR and α-H3K9me2 ChIP-qPCR data showed enrichment at a H3K9me2 reporter region (A) and depletion at a H3K4me3 reporter region (B) for two independent biological replicates. Control experiments with binding deficient TTD D142A mutant and IgG demonstrated the specificity of the assay. Control ChIP with α-H3K9me2 and CIDOPs with MPP8-CD and TAF3-PHD verified the assayed amplicons. For all regions *n* = 2, the bars represent the mean and error bars 0.95 confidence intervals (CI).

**B** Coordinates and amplicon size of H3K9me2 and H3K4me3 reporter regions used in the qPCR assays.

**C** CIDOP-seq and ChIP-seq data for the reporter loci used in qPCR assays. The amplicon borders are annotated with vertical bars in the middle of the window. All tracks in Reads Per Kilobase of transcript, per Million mapped reads (RPKM), y-axes start from 0. Coordinates in hg38.

**A**

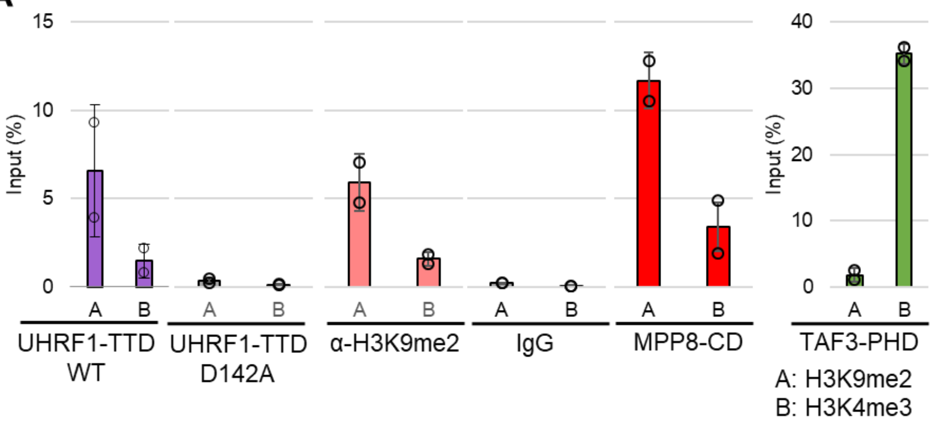


**B**

| Name | hg38 coordinates | Amplicon size (bp) |
|---|---|---|
| H3K9me2 | chr3:193713288-193713376 | 89 |
| H3K4me3 | chr7:104208109-104208217 | 109 |

**C**

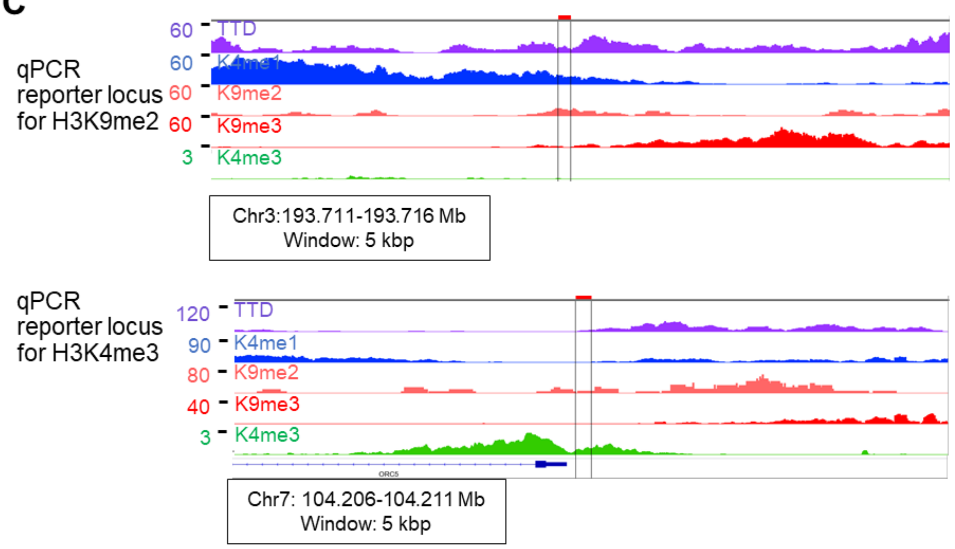

**Supplemental Figure 7. Additional browser views of the CIDOP-seq and ChIP-seq data shown in Figure 3C demonstrating reproducibility of experimental replicates.**

Data from hUHRF1-TTD CIDOP-seq and α-H3K9me2 ChIP-seq in two biological replicates showed good reproduction and were pooled. Representative browser views. All tracks in RPKM, y-axes start from 0. Coordinates in hg38.

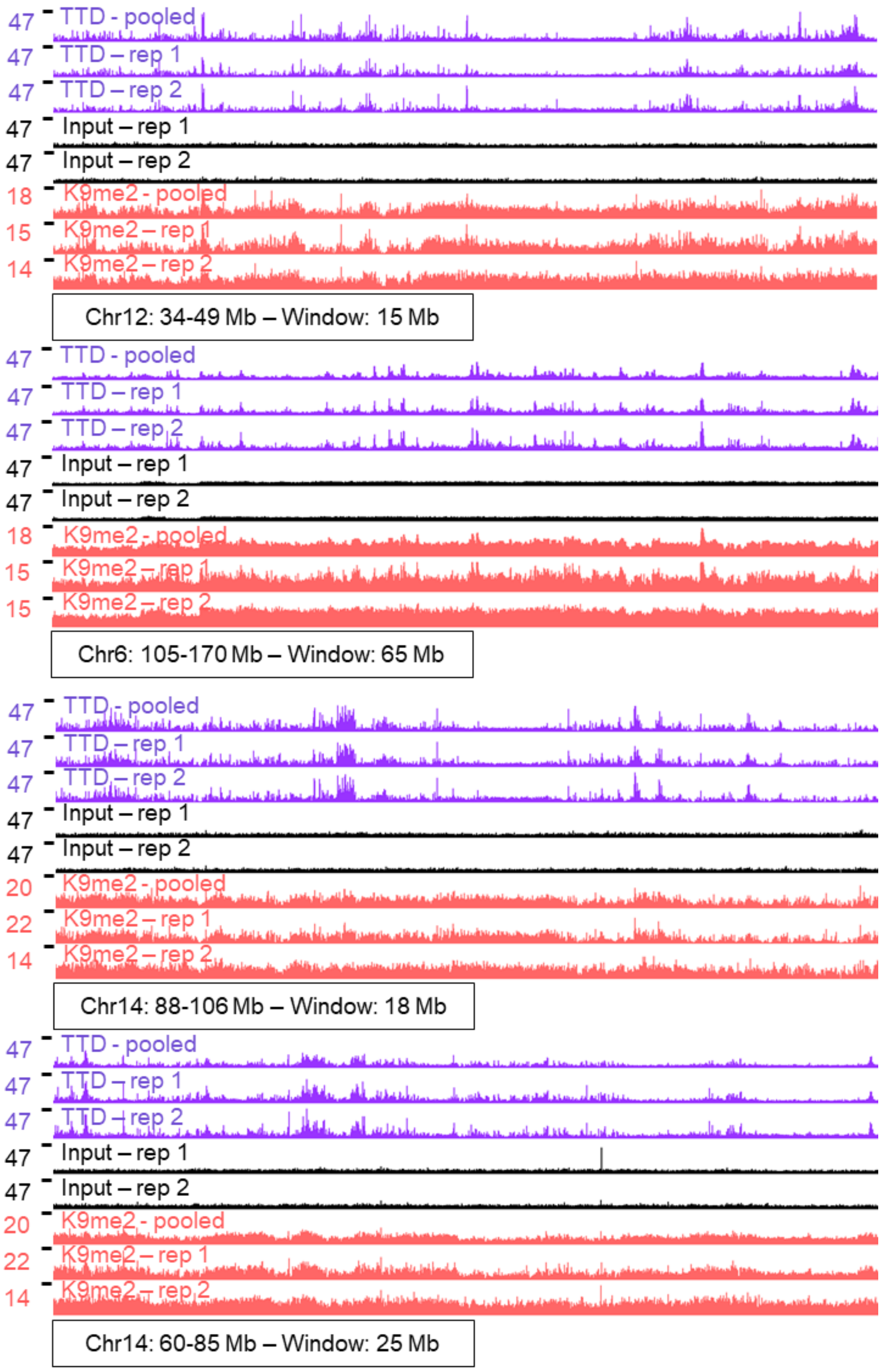


| Genome-wide *r* | | |
|---|---|---|
| hUHRF1- TTD | | |
| | rep. 1 | rep. 2 |
| Merged | 0.93 | 0.95 |

| Genome-wide *r* | | |
|---|---|---|
| α-H3K9me2 | | |
| | rep. 1 | rep. 2 |
| Merged | 0.93 | 0.86 |

**Supplemental Figure 8. Additional browser views of the CIDOP-seq and ChIP-seq data shown in Figure 3C demonstrating TTD CIDOP signal enrichment in regions with both H3K4me1 and H3K9me2/3.**

Additional representative data showing that hUHRF1-TTD CIDOP-seq is enriched in regions with both H3K4me1 and H3K9me2/3. All tracks in RPKM, y-axes start from 0. Coordinates in hg38.

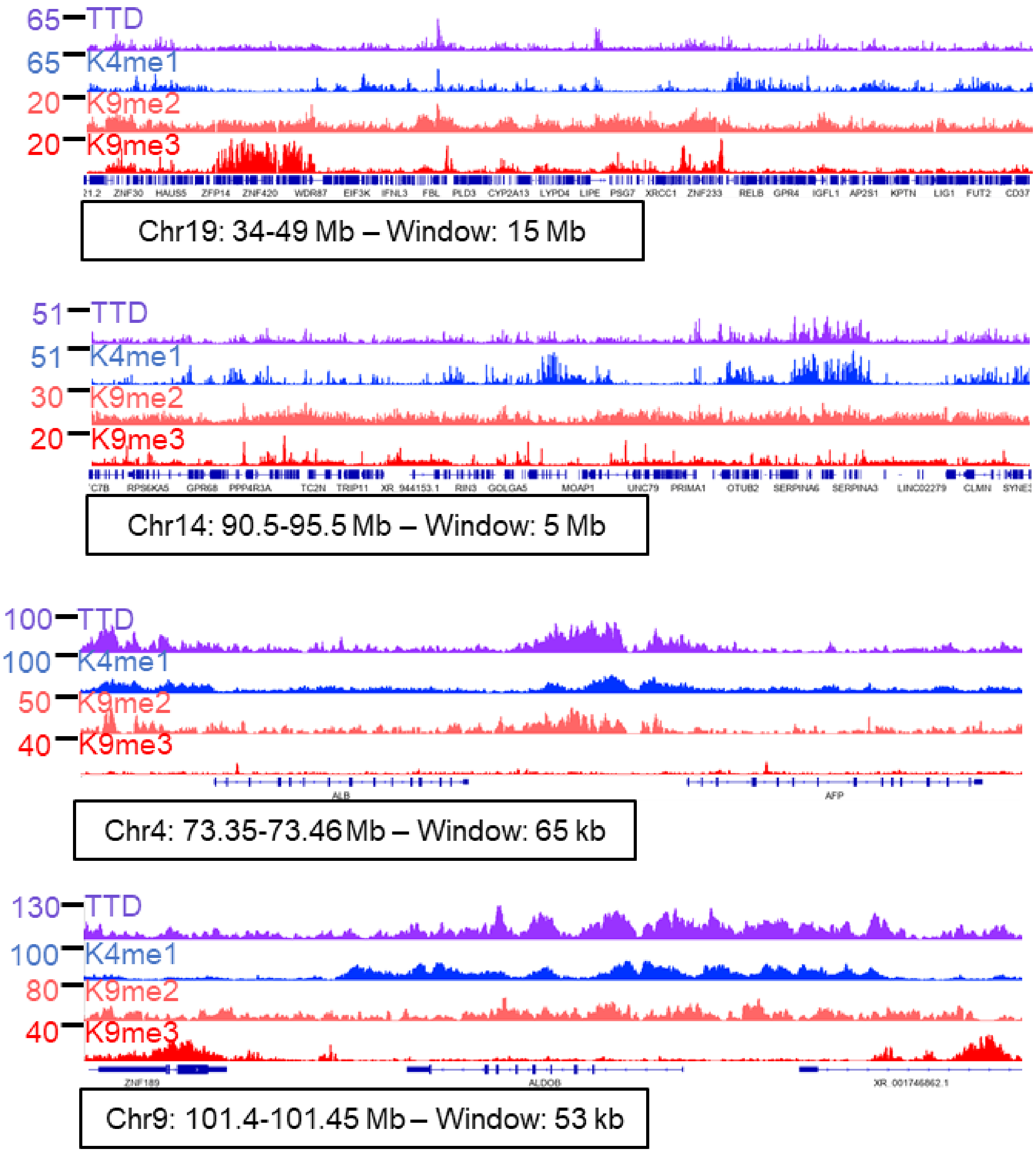

**Supplemental Figure 9. Additional browser views of the ChIP-seq data shown in Figure 3E demonstrating the broad genomic localisation of H3K9me2 and H3K9me3.**

H3K9me2 spans extremely wide regions of the genome. Chromosome arm-wide exemplary browser views showing megabase-wide regions of H3K9me2 enrichment. Data from a histone mark established as very broad (H3K9me3) show sharper and more defined regions of enrichment. HepG2 H3K9me3 data were taken from public datasets [2]. All tracks in RPKM, y-axes start from 0. All coordinates in hg38, gene annotation from RefSeq.

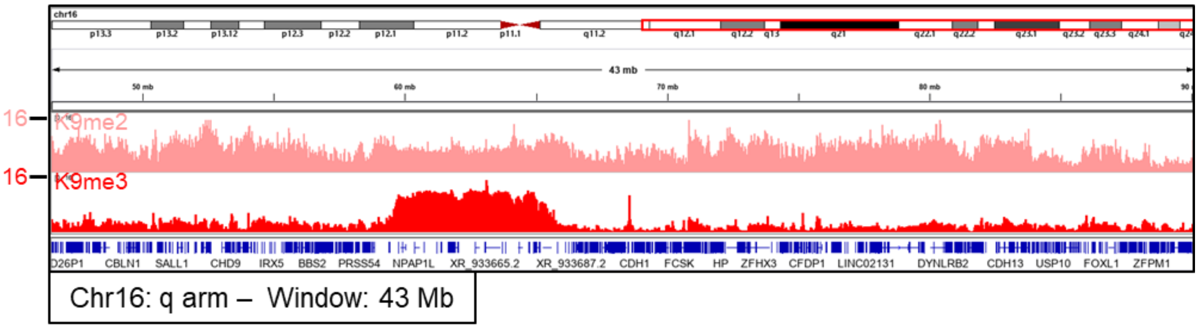


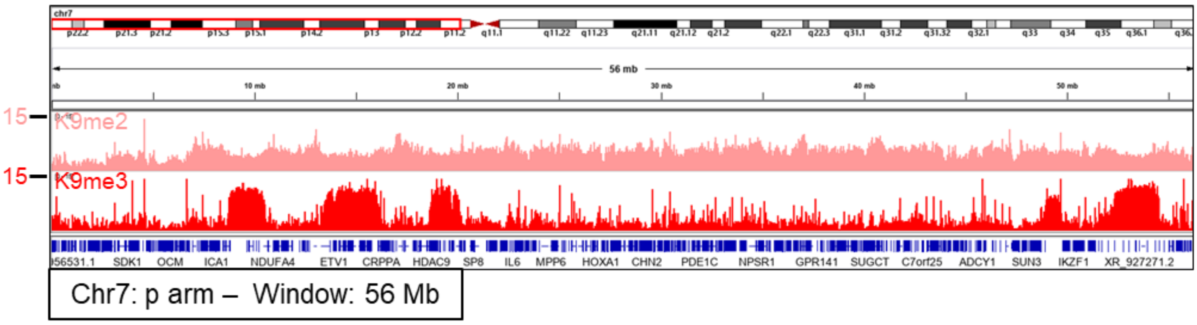

**Supplemental Figure 10. Additional data related to Figure 4.**

**A** The entire genome was divided into 1 kb bins, arranged by decreasing mean H3K9me2 signal, divided into deciles and the mean signal of each group was plotted. Central lines show the median, box borders are $25^{th}$ to $75^{th}$ percentile, and whiskers $5^{th}$ to $95^{th}$.

**B** TTD peaks overlap with H3K4me1 peaks. The number of overlaps refers to single counts and percentages refer to TTD peaks. Diagrams made using *Venn-Diagram-Plotter* (github.com/PNNL-Comp-Mass-Spec/Venn-Diagram-Plotter).

**C** Additional data related to Figure 4D. Additional H3K9me3 data are shown for the heatmap.

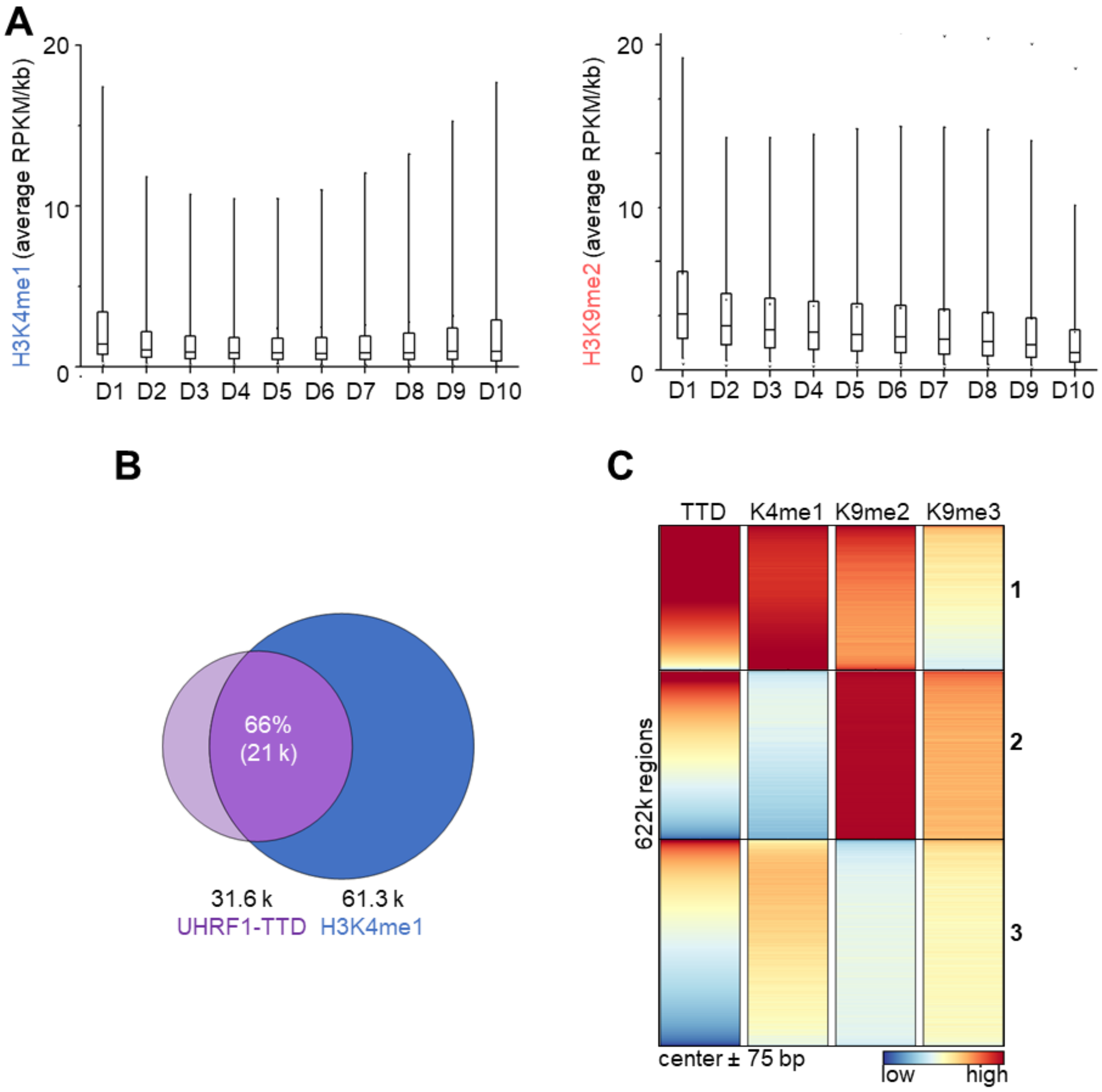

**Supplemental Figure 11. Additional data related to TTD binding to H3K4me1-K9me3 shown in Figure 4.**

**A** Heatmap of H3K9me3 peaks. Cluster 2 contains TTD signal with H3K4me1 and H3K9me3, in absence of H3K9me2. Heatmap of 86403 H3K9me3 peaks and their flanks, centered in the middle ± 2.5 kb, arranged by decreasing TTD signal.

**B** H3K4me1 peaks overlap 12% of the H3K9me3 peaks. The number of overlaps refers to single counts and the percentage refers to H3K4me1 peaks. Diagram made using *Venn-Diagram-Plotter* (github.com/PNNL-Comp-Mass-Spec/Venn-Diagram-Plotter).

**C** TTD can be recruited to H3K4me1 peaks that overlap H3K9me3 peaks, even without H3K9me2. In cluster 2 we see TTD signal with H3K4me1 and H3K9me3, in absence of H3K9me2. Heatmap of 4470 H3K4me1 peaks that overlap to ≥ 50% H3K9me3 peaks, centered in the middle ± 2.5 kb, clustered in 3 groups and arranged by decreasing H3K4me1 signal.

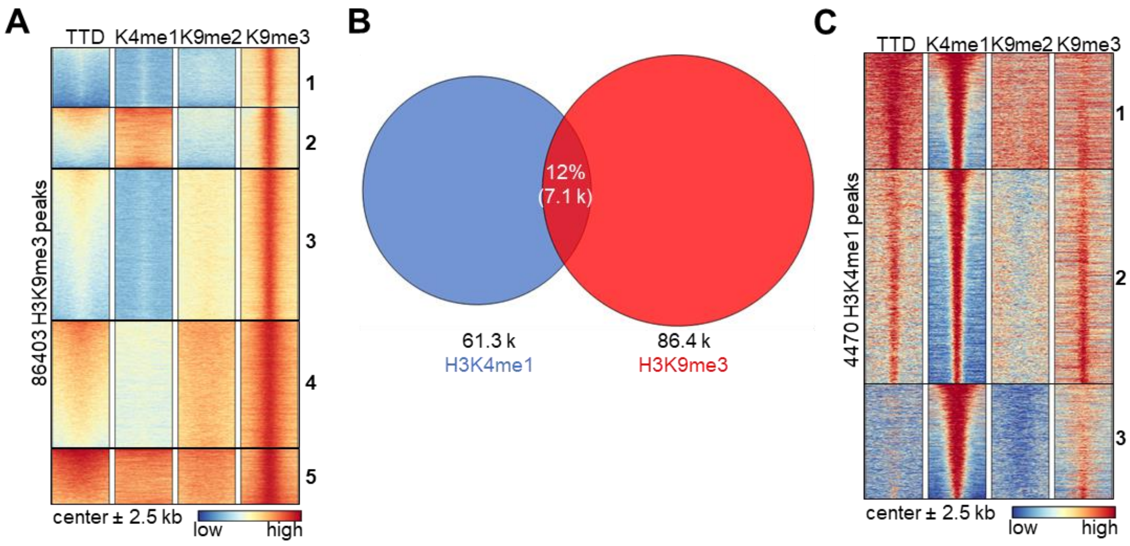

**Supplemental Figure 12. Additional data related to promoter and enhancer binding in HepG2 cells.**

**A** Genes robustly down-regulated in HepG2 vs. liver tissue (same genes as in Figure 5D) are HepG2 and liver specific, according to gene set enrichment analysis (GSEA) on *Enrichr* (maayanlab.cloud/Enrichr) [3].

**B** TTD peaks occur more frequently on enhancers, especially distal enhancers as annotated by *ChIP-Enrich* (chip-enrich.med.umich.edu).

**C** Control data for Whole Genome Bisulfite Sequencing (WGBS) signal on HepG2 enhancers. Heatmap of ~ 189k HepG2 enhancers identified by ChromHMM ± 2.5 kb (egg2.wustl.edu/roadmap) [4] centered in the middle and arranged by decreasing TTD signal.

**D** Genes with TTD peaks on their enhancers (found in HepG2, same genes as in Figure 6C) are HepG2 specific, according to GSEA on *Enrichr*.

**E** Exemplary browser views showing hUHRF1-TTD flanking peaks of selected cell-type specific transcription factors, ARID5B [5], FOXA1 aka HNF3α [6], HNF4A [6], and DNase [7] in HepG2 cells.

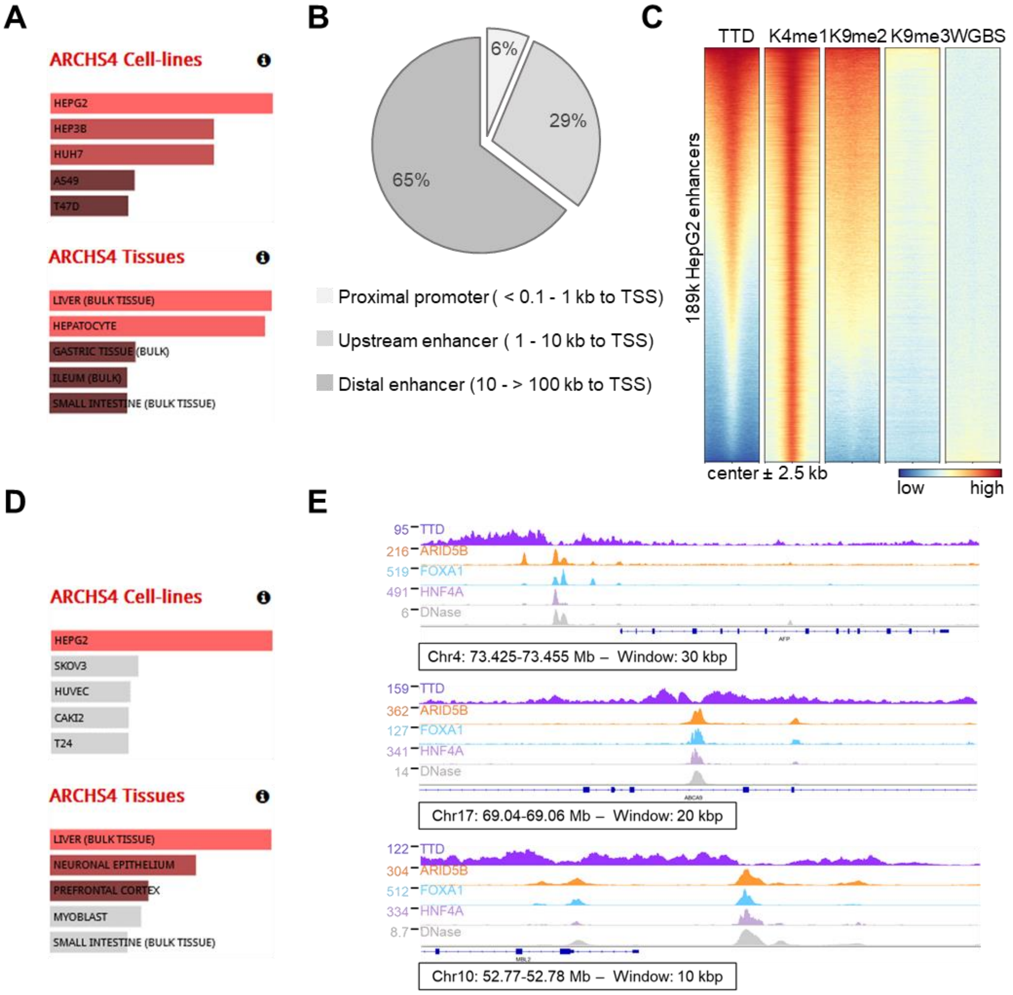

**Supplemental Figure 13. Additional data related to full-length mUHRF1 colocalization with H3K4me1 and gene regulation by UHRF1 in HCT116 cells shown in Figure 7.**

**A** Additional data referring to the mUHRF1 and H3K4me1 ChIP-seq shown in Figure 7B. Plot of the average mUHRF1 and H3K4me3 ChIP-seq signals in 2 kb bins genome-wide and Pearson´s correlation (*r*). Comparison with Figure 4G shows that mUHRF1 ChIP-seq correlates with H3K4me1 better than H3K4me3 in E14 mESC.

**B-D** Additional data related to Figure 7C-E.

**B** Counts of differentially regulated genes (DRGs) and corresponding enhancer regions using FC ≥ |1.5| from microarrays using UHRF1 KD treated HCT116 [8]. HCT116 cells with wild-type rescued cells over mock (WT) microarray had less DRGs than the Y188A rescued over mock (TTD*).

**C** TTD* up-regulated genes are enriched in H3K9me2 on their FANTOM5 enhancers [9]. Shown are the differentially regulated genes (DRGs) from the wild-type (WT) over mock non-responsive genes, sorted according to their status in Y188A mutant over mock (TTD*). The mean H3K9me2 signal of each group was plotted from HCT116 ChIP-seq data [10]. Central line is median, box borders are 25th to 75th percentile, and whiskers 5th to 95th. p values are from one-way ANOVA with Bonferroni correction. non-r. – non-responding.

**D** TTD* up-regulated genes are enriched in H3K4me1-K9me2 on their FANTOM5 enhancers [9]. Shown are the differentially regulated genes (DRGs) from all genes sorted according to their status in Y188A mutant over mock (TTD*). The mean H3K4me1 and H3K9me2 signal of each group was plotted from HCT116 ChIP-seq data [10, 11]. Central line is median, box borders are 25th to 75th percentile, and whiskers 5th to 95th. p values are from one-way ANOVA with Bonferroni correction. non-r. – non-responding.

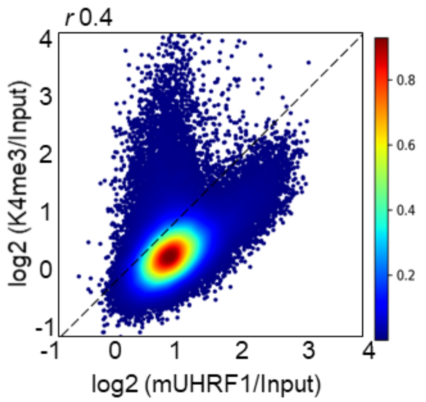


**B**

| WT vs Mock \|FC\| ≥ 1.5 | # genes | # enhancers |
|---|---|---|
| up-reg. | 267 | 1184 |
| down-reg. | 369 | 1284 |
| non-resp. | 20911 | 58798 |

| TTD* vs Mock \|FC\| ≥ 1.5 | # genes | # enhancers |
|---|---|---|
| up-reg. | 320 | 1420 |
| down-reg. | 451 | 1554 |
| non-resp. | 20776 | 58292 |

| WT non-resp. & TTD* vs Mock \|FC\| ≥ 1.5 | # genes | # enhancers |
|---|---|---|
| up-reg. | 115 | 414 |
| down-reg. | 152 | 501 |
| non-resp. | 20644 | 57833 |

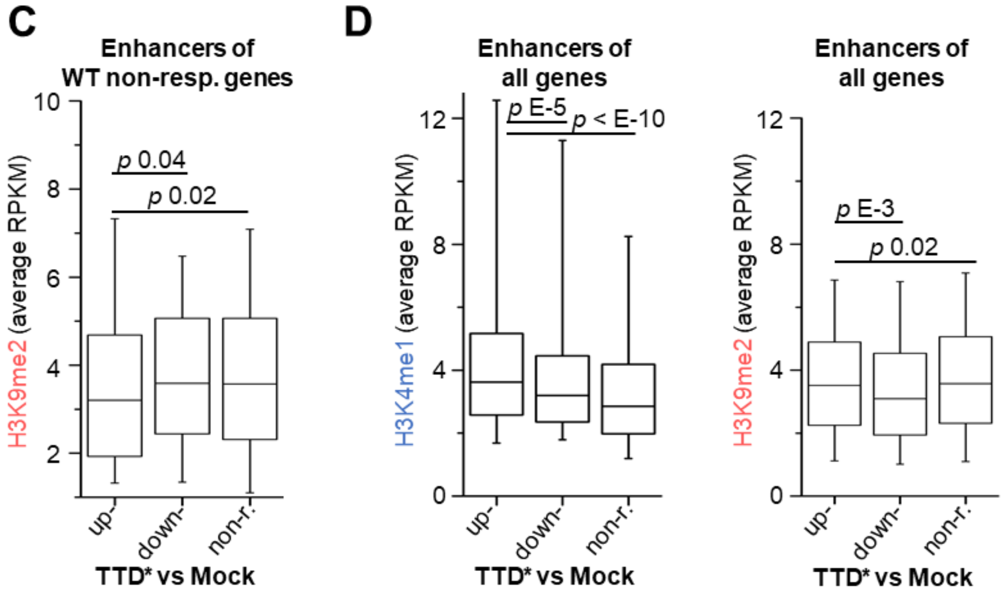

### Supplemental Tables

#### Supplemental Table 1. Oligonucleotides used for mutagenesis in this study.

| Description | Sequence |
|---|---|
| D142A | GAGTACGTCGccGCTCGGGAC |
| D142E | AATGAGTACGTCGAgGCTCGGGACACGAAC |
| D142N | GGTCAATGAGTACGTCaaCGCTCGGGACACGAAC |
| E153A | CGTGGTTTGctGCGCAGGTGG |
| E153D | GCGTGGTTTGAtGCGCAGGTGGTCAG |
| E153Q | GCGTGGTTTcAGGCGCAGGTGGTCAG |
| R207A | AGCGgccGCCCGCAC |
| R207L | CCGAGCGCtCGCCCGCAC |
| R207Q | AGCGCagGCCCGCAC |
| R207H | GTCCGAGCGCatGCCCGCAC |
| R207E | GTCCGAGCGgagGCCCGCAC |
| A208G | CGCGCGgCCGCACCAT |
| M224A | gaggtgggcCAGGTGGTCgcGCTCAACTACAAC |
| F278A | GTGGACGAAGTCgcgAAGATTTGACTC |

**Supplemental Table 2. Fluorescent histone H3 peptides used in this study.** FITC – fluorescein isothiocyanate; F – fluorescein

| Abbreviation | Residues | Sequence |
|---|---|---|
| H3(1-17) | 1-17 | ARTKQTARKSTGGKAPR-K(FITC) |
| H3K4me1 | 1-17 | ARTK(me1)QTARKSTGGKAPR-K(FITC) |
| H3K9me2 | 1-17 | ARTKQTARK(me2)STGGKAPR-K(FITC) |
| H3K9me3 | 1-19 | ARTKQTARK(me3)STGGKAPRKQ-K(FITC) |
| H3K4me1-K9me2 | 1-19 | ARTK(me1)QTARK(me2)STGGKAPRKQ-K(FITC) |
| H3K4me1-K9me3 | 1-19 | ARTK(me1)QTARK(me3)STGGKAPRKQ-K(F) |
| H3R8me2s-K9me3 | 1-19 | ARTKQTAR(me2s)K(me3)STGGKAPRKQ-K(F) |

**Supplemental Table 3. Antibodies used for ChIP and/or western blots, with conditions of use.**

| | α-H3K9me2 | IgG ms | α-H3K4me1 | IgG rb | α-H3K4me3 |
|---|---|---|---|---|---|
| Company | abcam | Invitrogen | abcam | R&D System | abcam |
| Cat. no. | ab1220 | 31903 | ab176877 | AB-105-C | ab8580 |
| RRID | AB_449854 | AB_10959891 | AB_2637011 | AB_354266 | AB_306649 |
| Lot no. | GR3377057-1 | WE3290156 | GR208955-11 | ER1619081 | GR3362382-2 |
| Dilution (v/v) | 1:1000 | not applicable | 1:5000 | n.a. | 1:2000 |
| 2nd lot no. | - | - | - | | GR3190162-2 |

**Supplemental Table 4. Reagents and conditions used for CIDOP and ChIP from HepG2 mononucleosomes.**

| | MPP8-CD | TAF3-PHD | UHRF1-TTD WT | UHRF1-TTD D142A | α-H3K9me2 (ab1220) | IgG ms (31903) | α-H3K4me1 (ab176877) | IgG rb (AB-105-C) |
|---|---|---|---|---|---|---|---|---|
| $c_{final}$ | 0.5 µM | 2 µM | 2 µM | 2 µM | 2 µg per ChIP | 2 µg per ChIP | 4 µg per ChIP | 4 µg per ChIP |
| nmol | 0.25 | 1.0 | 1.0 | 1.0 | n.a. | n.a. | n.a. | n.a. |
| Bead V (µl) | 12 | 12 | 12 | 12 | 20 | 20 | 20 | 20 |
| Chromatin (nmol) | 300 | 300 | 300 | 300 | 150 | 150 | 300 | 300 |
| Chromatin (µg) | 60 | 60 | 60 | 60 | 30 | 30 | 60 | 60 |
| Wash steps | PB200 | PB200 | PB200 | PB200 | PB200 | PB200 | PB200 | PB200 |
| Rinse steps | T | T | T | T | TLi | TLi | T | T |

**Supplemental Table 5. Oligonucleotides used for qPCR assays in this study.**

| | |
|---|---|
| qPCR H3K9me2 fwd | ATGATTATGAGCCCACCAGGC |
| qPCR H3K9me2 rvs | AGAGTCAGCCTTTGATGCCA |
| qPCR H3K4me3 fwd | ACTCTCTTCTCGCTGGTCCT |
| qPCR H3K4me3 rvs | TCCATGTCGTCTCCTTAGCC |

**Supplemental Table 6. NGS public datasets used in this study.** Published ChIP-seq data were downloaded as fastq files or pre-processed ENCODE data (DNase, WGBS). Microarray data were downloaded as pre-processed txt files containing the Lowess normalized $log_2$(FC) ratio.

| Cells | Name | GEO code | Sample codes | Antibody | Ref. |
|---|---|---|---|---|---|
| HepG2 | H3K4me1 | GSM3019940 | SRX3733792 SRR6761496 | Diagenode, C15410194, lot A1863-001D | [2] |
| HepG2 | H3K9me3 | GSM3019942 | SRX3733794 SRR6761498 | Diagenode, C15410193, lot A1671-001P | [2] |
| HepG2 | Input | GSM3019946 | SRX3733798 SRR6761502 | n.a. | [2] |
| HepG2 | FOXA1 | GSM803461 | SRX100506 SRR351749-50 | Santa Cruz, sc-6553 | [6] |
| HepG2 | ARID5B | GSM2797486 | SRX3230377 SRR6117651-2 | α-FLAG | [5] |
| HepG2 | HNF4A | GSM803460 | SRX100505 SRR351747-8 | Santa Cruz, sc-8987 | [6] |
| HepG2 | DNase | GSE90300 | ENCFF113VII ENCFF546MZK | n.a. | [7] |
| HepG2 | WGBS | GSE86764 | ENCSR881XOU ENCFF601FCO | n.a. | [7] |
| HCT116 | H3K4me1 | GSM2058064 | SRX1568641 SRR3157819 | Abcam, ab8895 | [11] |
| HCT116 | H3K9me2 | GSM2916019 | SRX3566210 SRR6476346 | Abcam, ab1220 | [10] |
| HCT116 | WT | GSM3355169 | n.a. | n.a. - microarray | [8] |
| HCT116 | TTD* | GSM3355170 | n.a. | n.a. - microarray | [8] |
| E14 mESC | H3K4me3 | GSM3123475 | SRX4017932 SRR7088938 | Millipore, 07-473 | [12] |
| E14 mESC | mUHRF1 | GSM3123487 | SRX4017944 SRR7088950 | Santa Cruz, 98817 | [12] |
| E14 mESC | input | GSM3123488 | SRX4017945 SRR7088951 | n.a. | [12] |
| E14 mESC | H3K4me1 | GSM1847705 | SRX1141888 SRR2154876 | Abcam, ab8895 | [13] |

**Supplemental data provided as Excel files**

**Supplemental File 1. UHRF1-TTD and H3 PTMs overlaps in ChIP-Atlas database.** Raw and processed output of peak overlap analysis from the ChIP-Atlas database (chip-atlas.org) used to construct the diagram shown in Figure 4E.

**Supplemental File 2. ChIP-Enrich results for UHRF1-TTD enriched refTSS-cluster 1.** ChIP-Enrich (chip-enrich.med.umich.edu) output and Revigo (revigo.irb.hr) input used to construct the network shown in Supplemental Figure 8A.

**Supplemental File 3. TSSs of genes with ≥ 2-fold change in expression.** Coordinates from refTSS and pTPM values from Human Protein Atlas (proteinatlas.org) used to construct the heatmap shown in Figure 5C.

**Supplemental File 4. UHRF1-TTD peaks analyzed by ChIP-Enrich-hybrid.** ChIP-Enrich (chip-enrich.med.umich.edu) output and Revigo (revigo.irb.hr) input used to construct the network shown in Supplemental Figure 8C.

**Supplemental File 5. UHRF1-TTD and TF overlaps in ChIP-Atlas database.** Raw and processed output of peak overlap analysis from the ChIP-Atlas database (chip-atlas.org) used to construct the diagram shown in Figure 6D.
